## Supplementary Figures and Tables for "Spatially Aware Dimension Reduction for Spatial Transcriptomics"

Figure S1

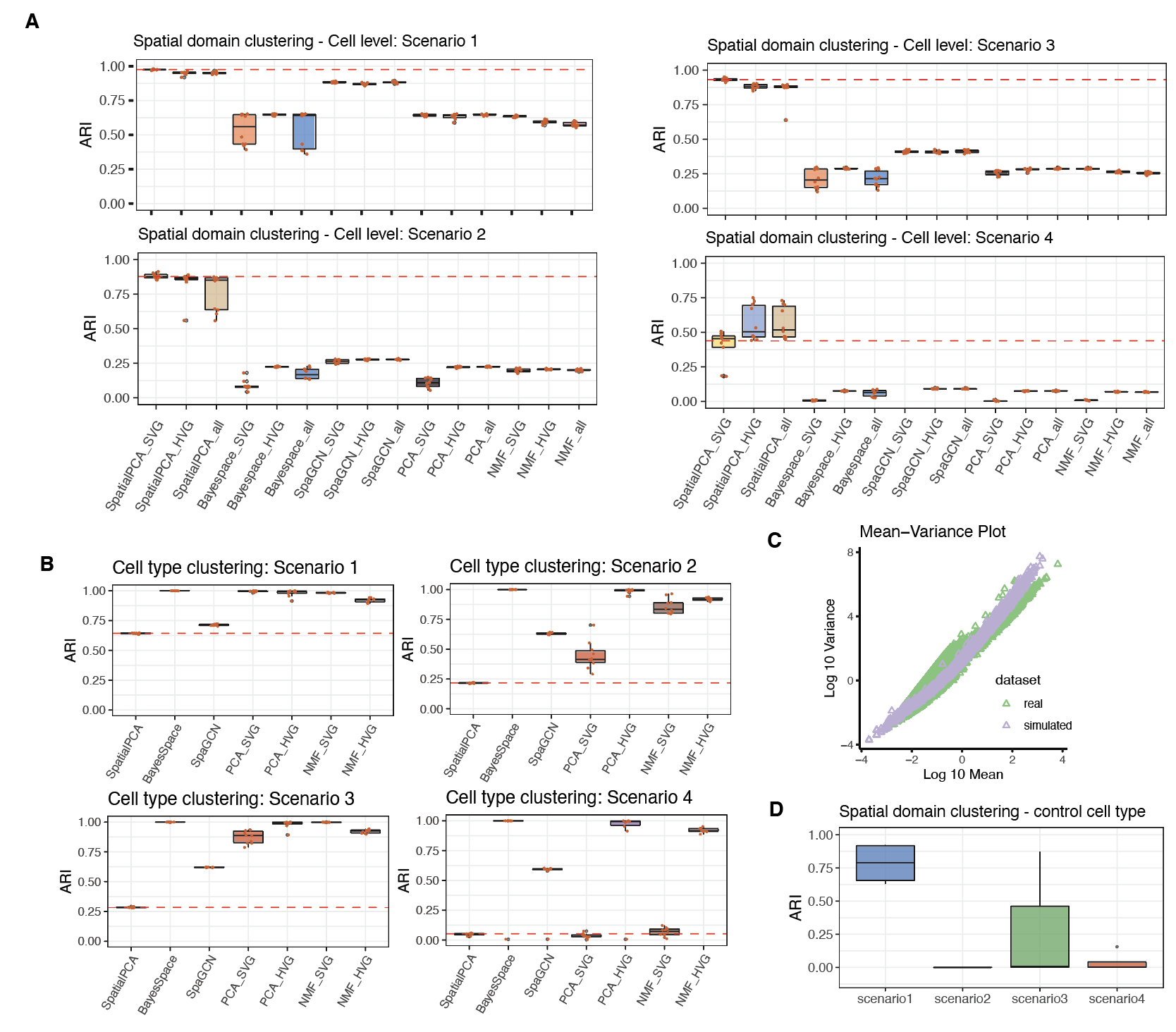

**Figure S1. Simulation results for spatial domain clustering and cell type clustering in single cell resolution.** (**A**) Spatial domain clustering results of different methods paired with spatially variable genes (SVGs), highly variable genes (HVGs) and all genes in four simulation scenarios. (**B**) Cell type clustering results of different methods. SpatialPCA has highest ARI in spatial domain clustering and lowest ARI in cell type clustering, highlights the different goals in spatial domain and cell type detection. In SpatialPCA we aim to identify spatial domains. (**C**) The mean and variance relationship between real scRNA-seq count data and simulated single cell count data are consistent. (**D**) After controlling for cell types as covariates in SpatialPCA, the clustering performance is the best when there is one dominant cell type in each spatial domain (scenario 1), and the performance reduces when there are multiple cell types mixed in each spatial domain (scenario 2, 3 and 4). These results highlight the spatial domains are driven by the cell type compositions.

Figure S2

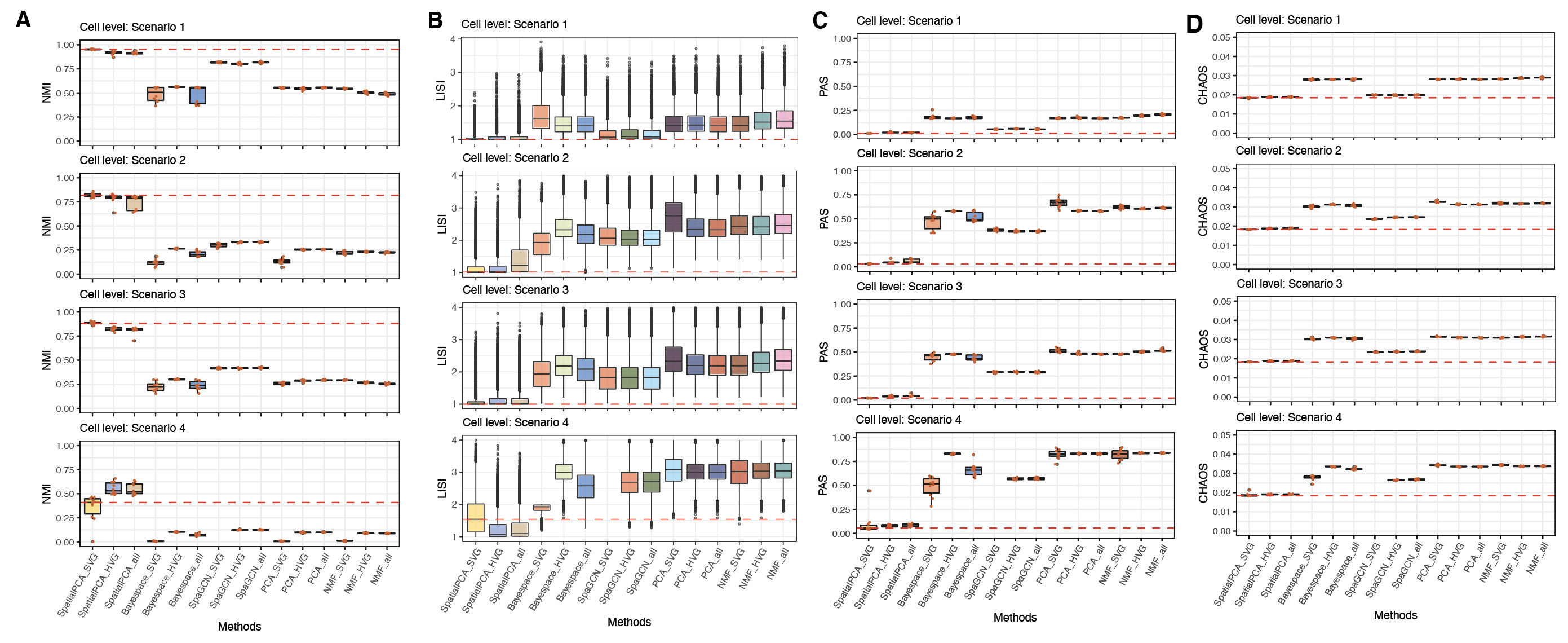

**Figure S2. Simulation results for spatial domain clustering in single cell resolution.** Spatial domain clustering results of different methods paired with SVGs, HVGs and all genes in four simulation scenarios, in terms of (**A**) NMI (the higher the better), (**B**) LISI (the lower the better), (**C**) PAS (the lower the better), and (**D**) CHAOS (the lower the better) scores.

Figure S3

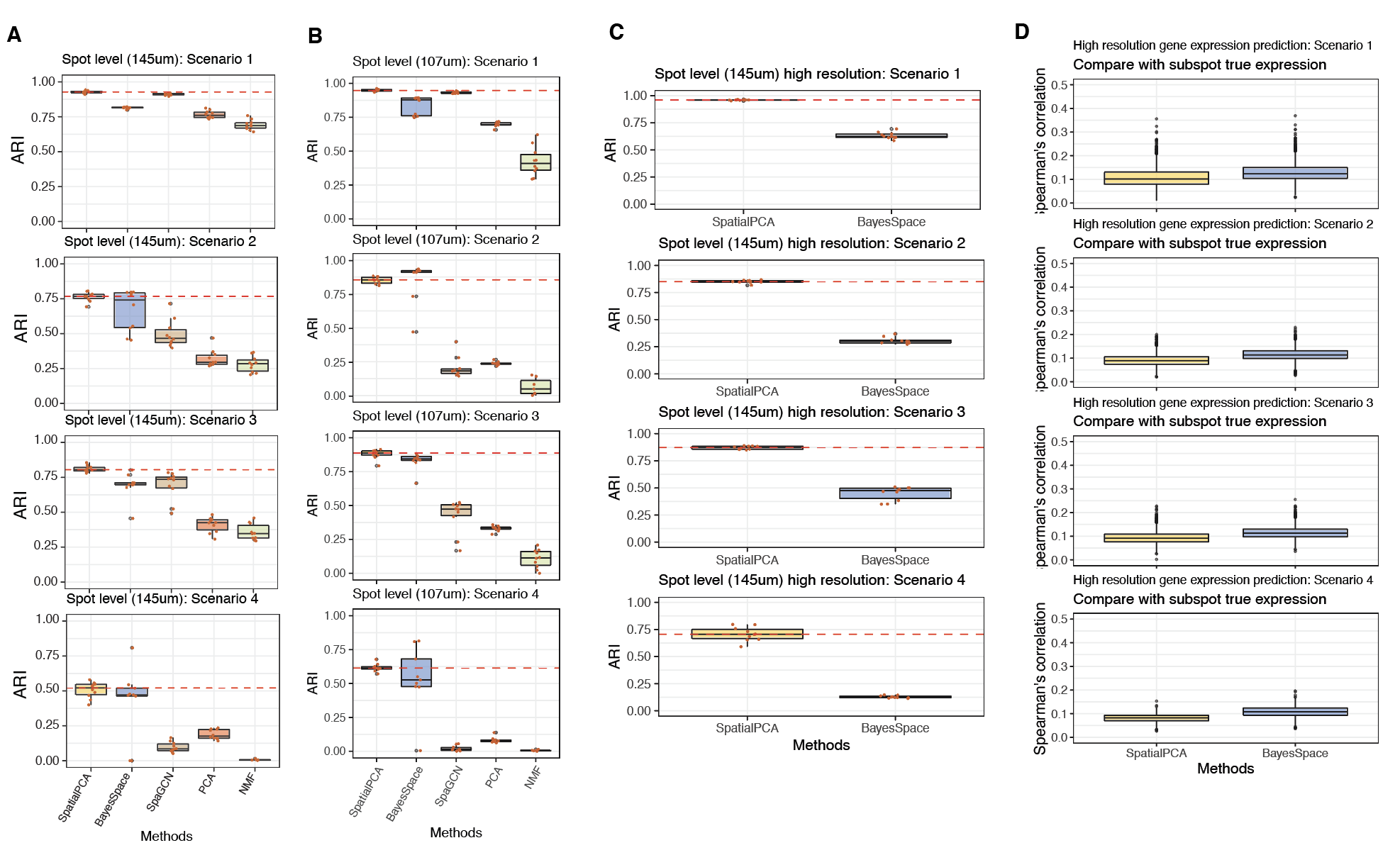

**Figure S3.** **Simulation results for spatial domain clustering at spot level.** (**A**) Spatial domain clustering results of different methods at spot diameter being 145um. (**B**) Spatial domain clustering results of different methods at spot diameter being 107um. (**C**) High resolution spatial map clustering results for spot level simulation at spot diameter being 145um. We compared SpatialPCA with BayesSpace. (**D**) High resolution gene expression prediction results for spot level simulation at spot diameter being 145um. We compared SpatialPCA with BayesSpace.

Figure S4

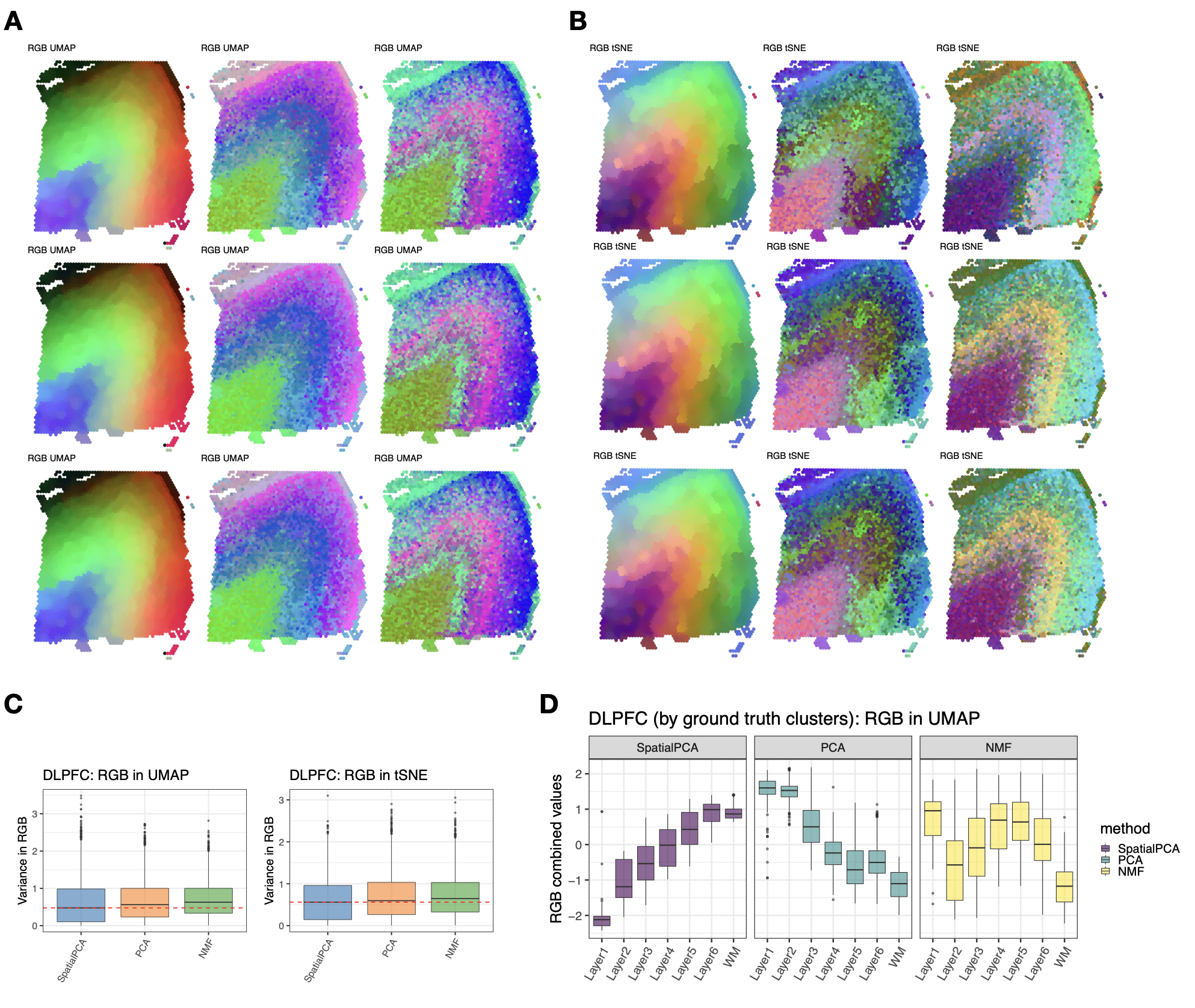

**Figure S4.** **RGB plots for the DLPFC data**. (**A-B**) For SpatialPCA, PCA, and NMF, we summarized the inferred low dimensional components into three UMAP (left panels) or tSNE components (right panels) and visualized the three resulting components with red/green/blue (RGB) colors through the RGB plot. The RGB plot from SpatialPCA displays laminar organization of the cortex and show less color differences within a local area. We also scaled up spatial PCs/regular PCs 10 times (second row) and 20 times (third row) to see the influence of range of the PCs to RGB plots, the tSNE/UMAP results and RGB plots have very similar patterns as shown at the original scale (first row). (**C**) We found in SpatialPCA, the weighted RGB values have lower variance than PCA or NMF in nearby spots. (**D**) The RGB plots in SpatialPCA show a smoother transition of colors between adjacent cortical layers. The cortical layers are labeled based on ground truth annotations.

Figure S5

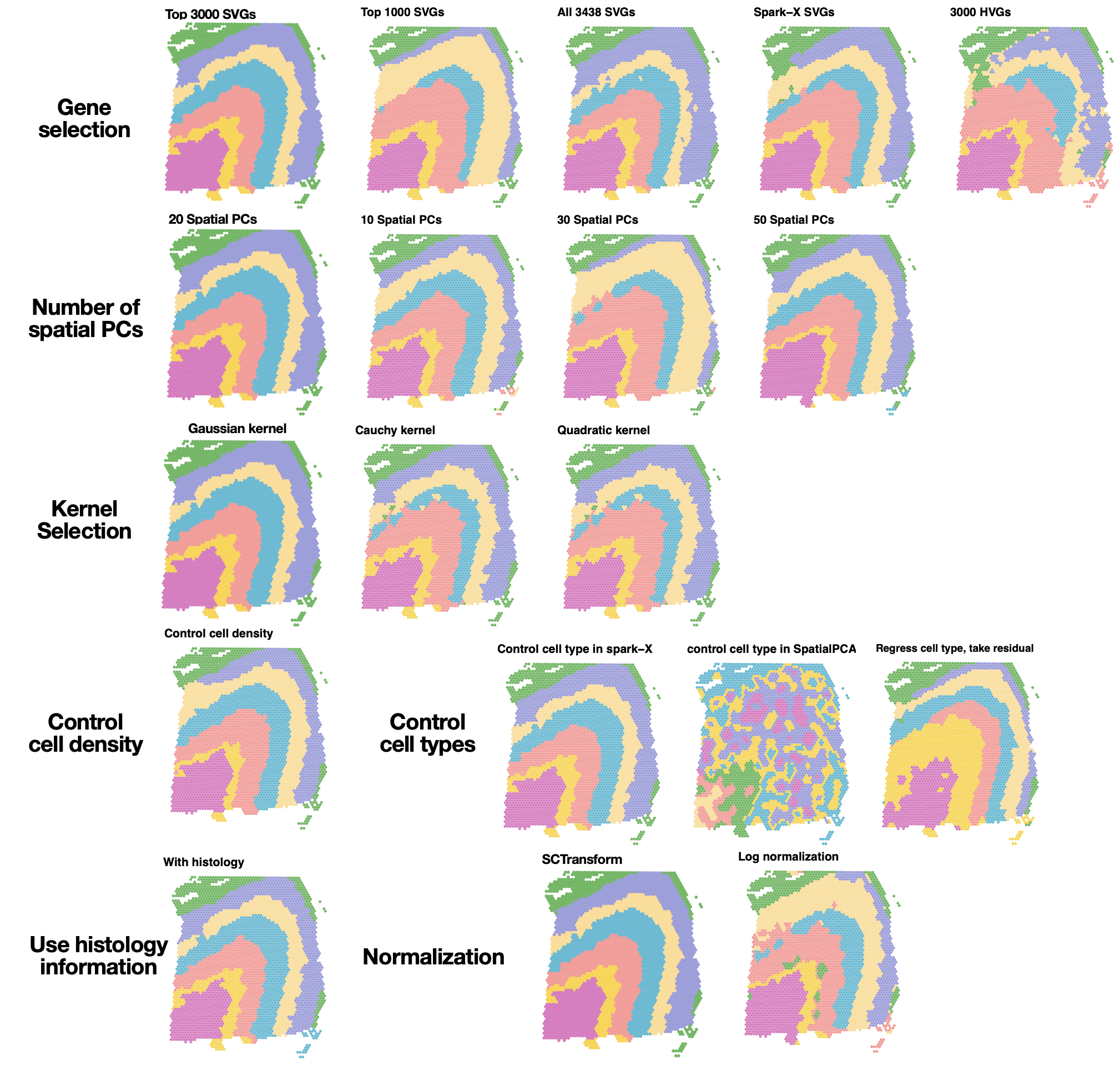

**Figure S5. Sensitivity analysis in the DLPFC data**. **First row**: clustering results obtained using a different set of input genes, include top 3000 SVGs, top 1000 SVGs, all SVGs detected by SPARK, all SVGs detected by SPARK-X, and top 3000 HVGs. In SPARK and SPARK-X, the adjusted p value cut-off for determining the SVGs is set to be 0.05. **Second row**: clustering results obtained using either the top 20, 10, 30, or 50 spatial PCs. **Third row**: clustering results based on spatial PCs extracted using either the Gaussian kernel, the Cauchy kernel, or the rational quadric kernel. **Fourth row**: Left: clustering results obtained by controlling for cell density in the spots. The cell density information of the DLPFC data was obtained from their original paper. Right: clustering results obtained by controlling cell types when selecting SVGs in SPARK-X; controlling cell types in SpatialPCA; controlling cell types by regressing them out from the input gene expression and take the residuals. **Fifth row**: Left: clustering results obtained by taking the histology information as a third dimension in location matrix. The histology information was extracted from the H&E image following SpaGCN. Right: clustering results obtained by SpatialPCA with gene expression normalized through SCTransform normalization or log normalization.

Figure S6

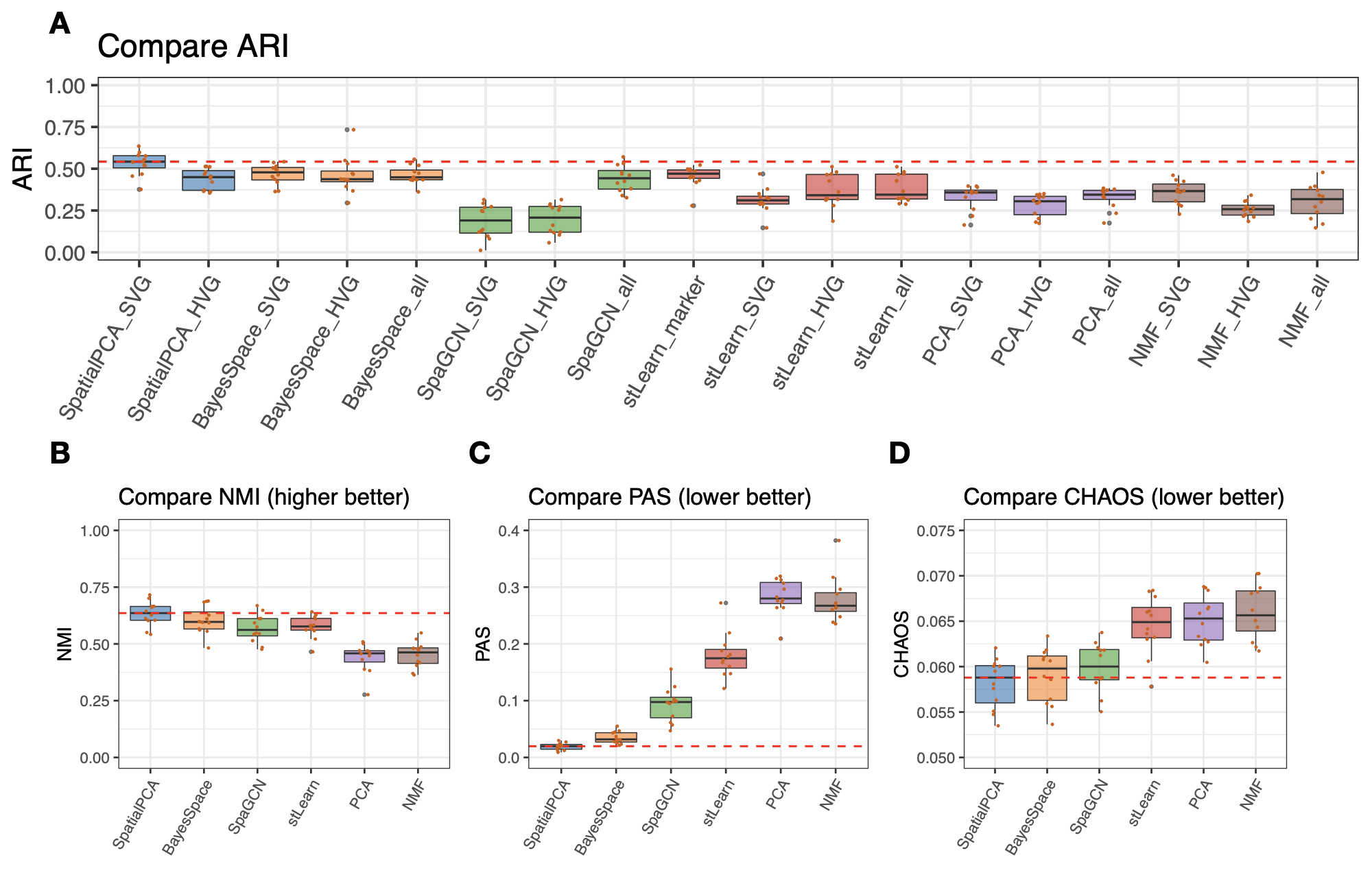

**Figure S6. Clustering results obtained based on different methods in the DLPFC data.** (**A**) Clustering results measured by ARI (the higher the better). In dimension reduction methods (SpatialPCA, PCA, and NMF), clustering was performed based on the inferred low-dimensional components. For spatial domain clustering methods (BayesSpace and SpaGCN), clustering was performed based the default settings. All the methods are paired with SVGs, HVGs and all genes. (**B**-**D**) Clustering results measured by NMI (the higher the better), PAS (the lower the better), and CHAOS (the lower the better) in different methods with their default settings. Clustering results of PCA and NMF are obtained with SVGs.

Figure S7

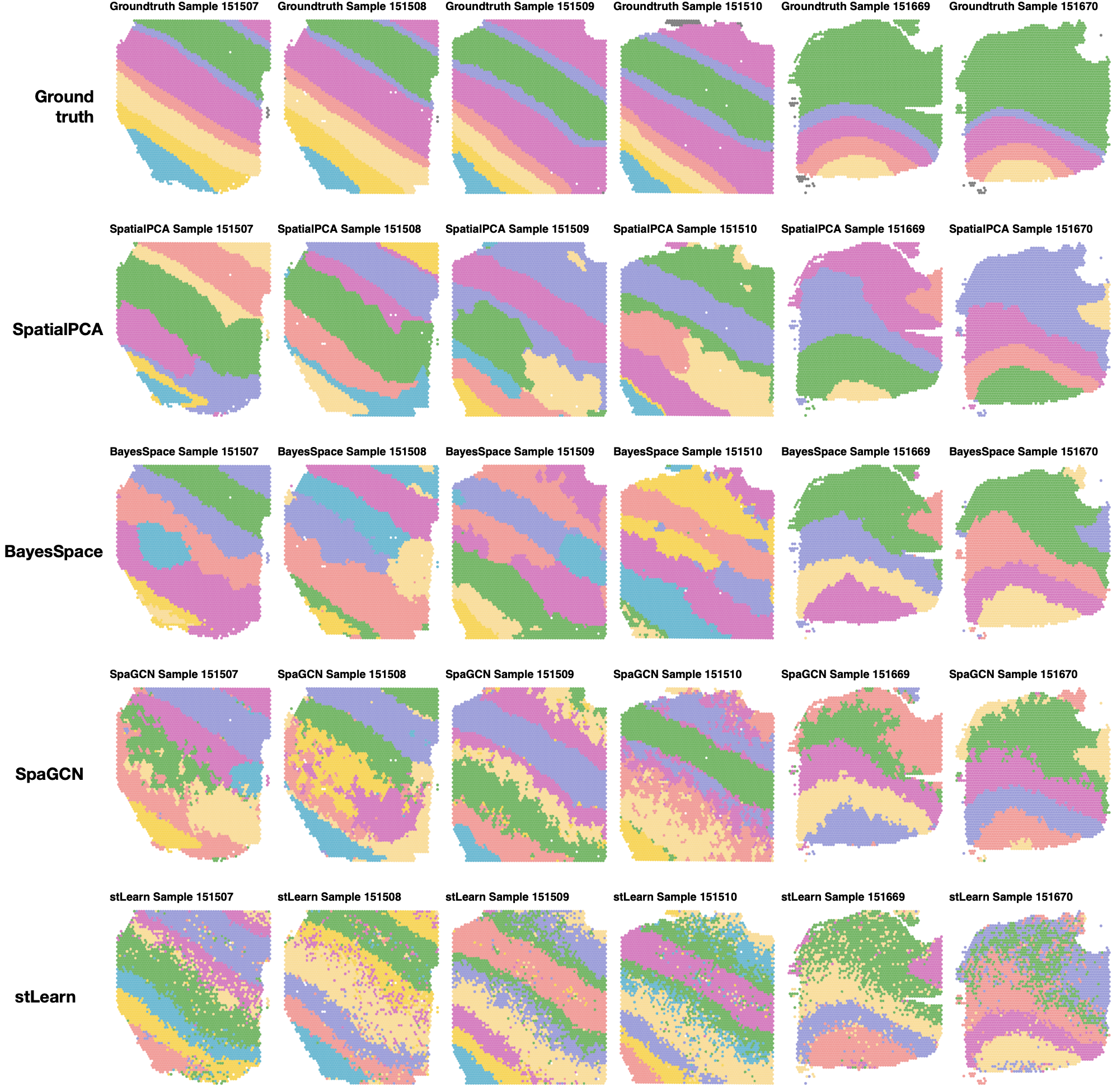

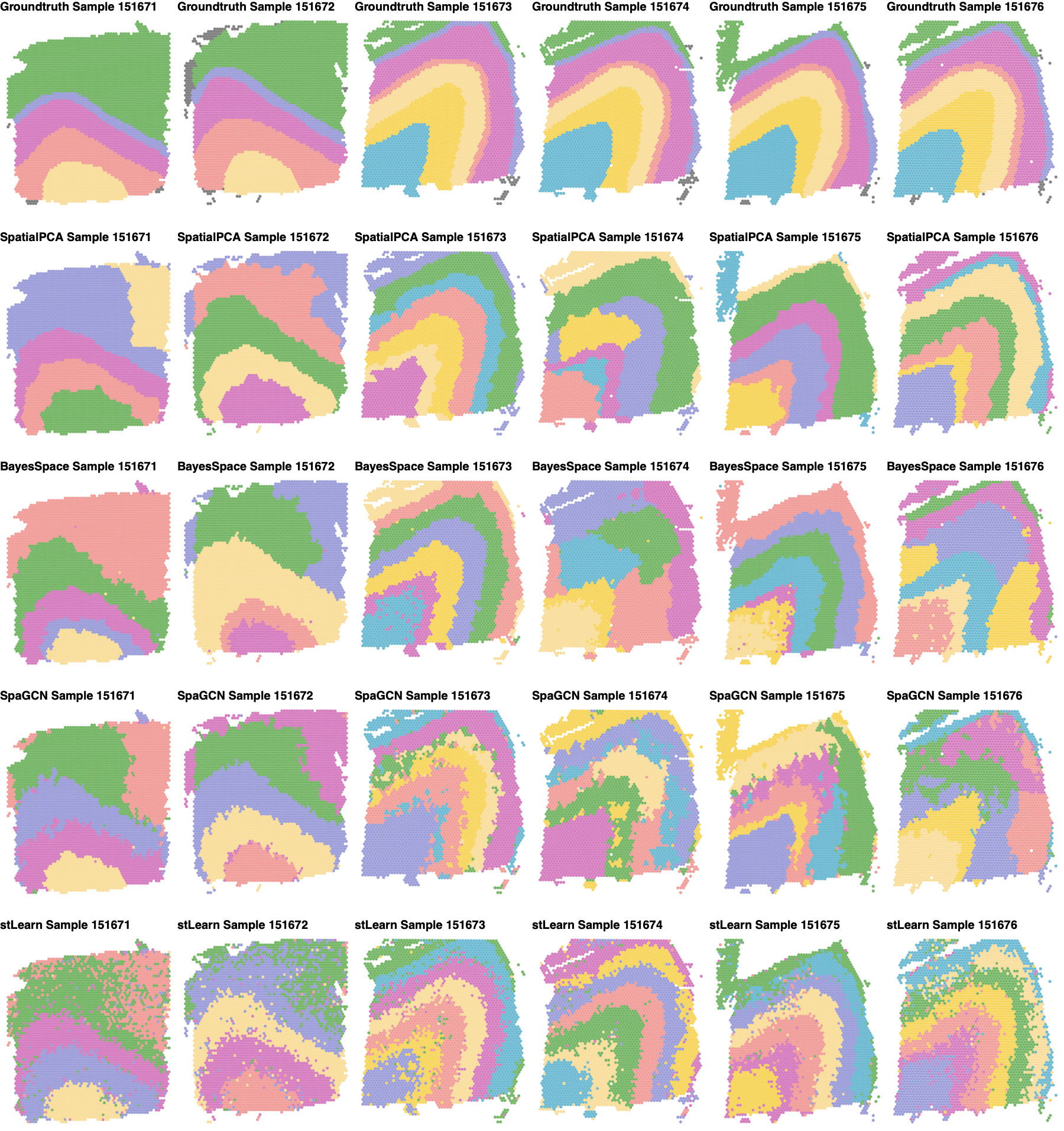

**Figure S7. Clustering results of different spatial domain detection methods in DLPFC across all twelve samples.** **First row**: Ground truth annotation of all twelve samples**. Second row**: SpatialPCA clustering results. **Third row**: BayesSpace clustering results. **Fourth row**: SpaGCN clustering results. **Fifth row**: stLearn clustering results.

Figure S8

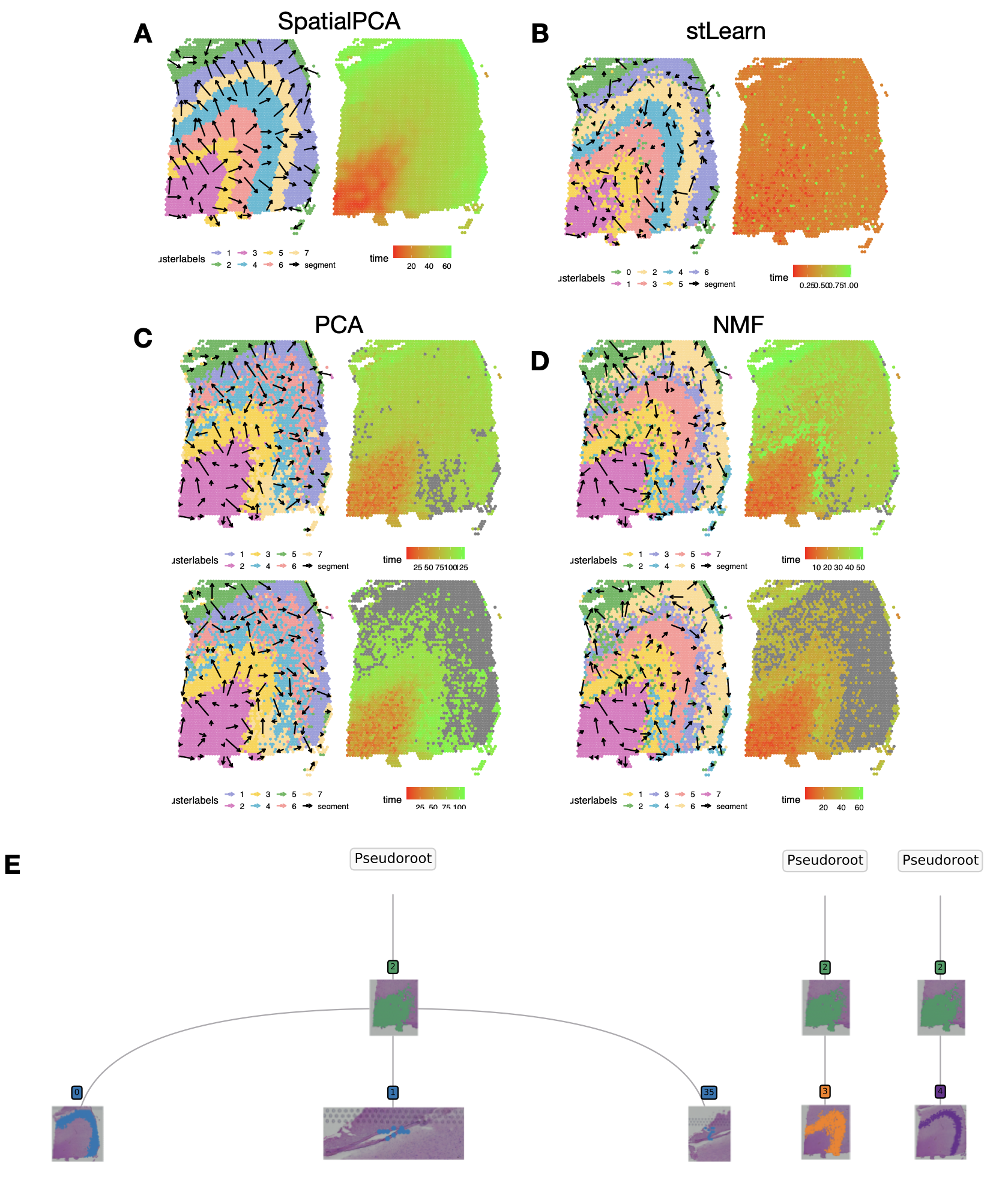

**Figure S8.** **Spatial trajectory inference results in the DLPFC data**. **(A)**: Visualizaton of the trajectory inferred by SpatialPCA. Left: Arrows point from tissue locations with low pseudo-time to tissue locations with high pseudo-time. Color represents different tissue regions. Right: Visualization of pseudotime inferred from spatial PCs in SpatialPCA. (**B**): Visualizaton of the pseudotime inferred by stLearn. We plotted the arrows in the same way as in SpatialPCA. (**C**-**D**): Visualizaton of the trajectories inferred from PCs in PCA and NMF. (**E**): Visualization of trajectories inferred by stLearn. The stLearn considers a pair of clusters at each time and find the order between clusters.

Figure S9

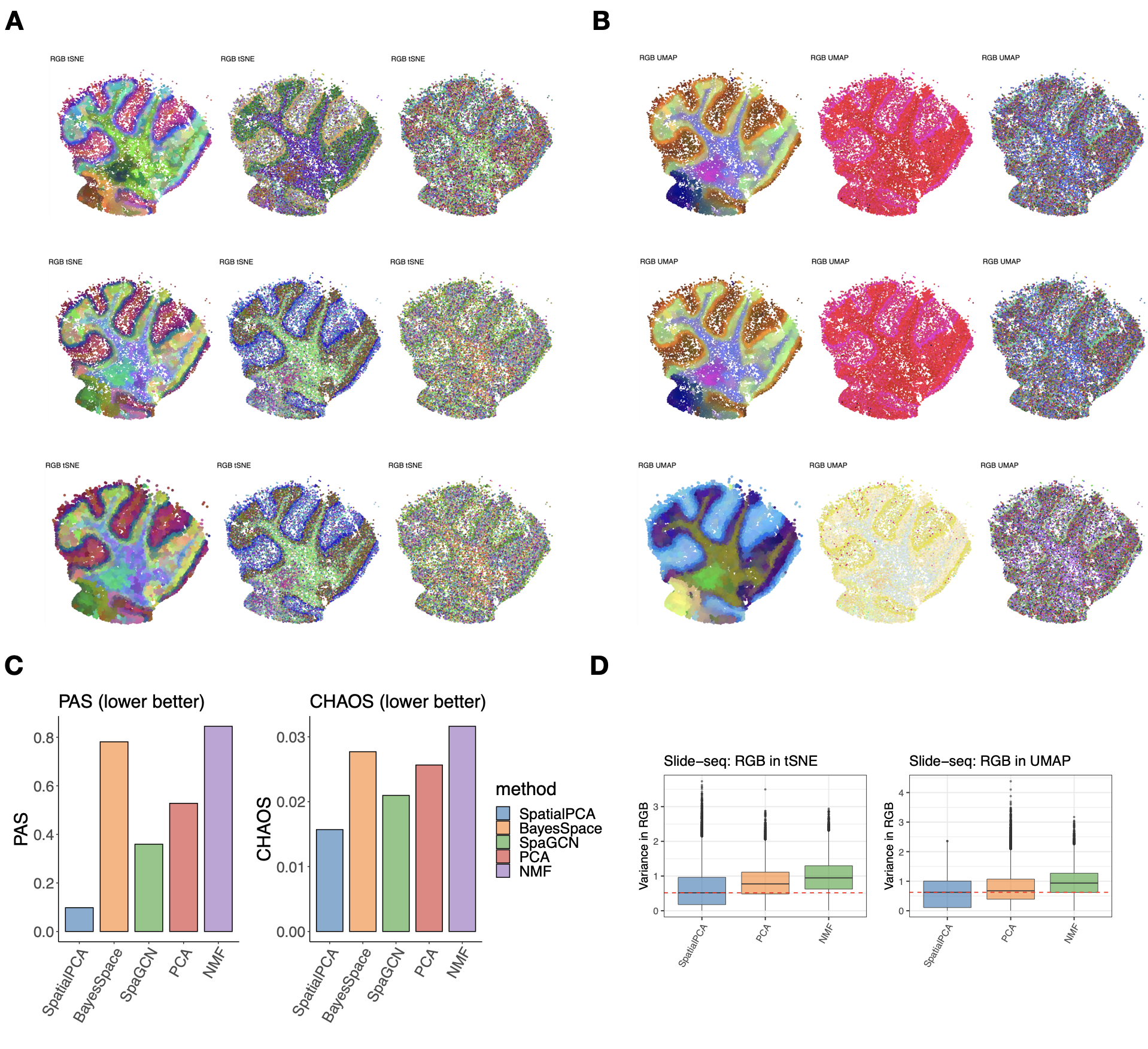

**Figure S9.** **RGB plots for the Slide-seq data**. (**A-B**) For SpatialPCA, PCA, and NMF, we summarized the inferred low dimensional components into three UMAP (right panels) or tSNE components (left panels) and visualized the three resulting components with red/green/blue (RGB) colors through the RGB plot. The RGB plot from SpatialPCA displays tissue structure organization of the cerebellum and show less color differences within a local area. We also scaled up spatial PCs/regular PCs 10 times (second row) and 20 times (third row) to see the influence of range of the PCs to RGB plots, the tSNE/UMAP results and RGB plots have very similar patterns as shown at the original scale (first row). (**C**) The PAS and CHAOS scores (the lower the better) of clustering results in SpatialPCA is the lowest compared with BayesSpace, SpaGCN, PCA and NMF. (**D**) The weighted RGB values in SpatialPCA have lower variance than PCA or NMF in nearby spots.

Figure S10

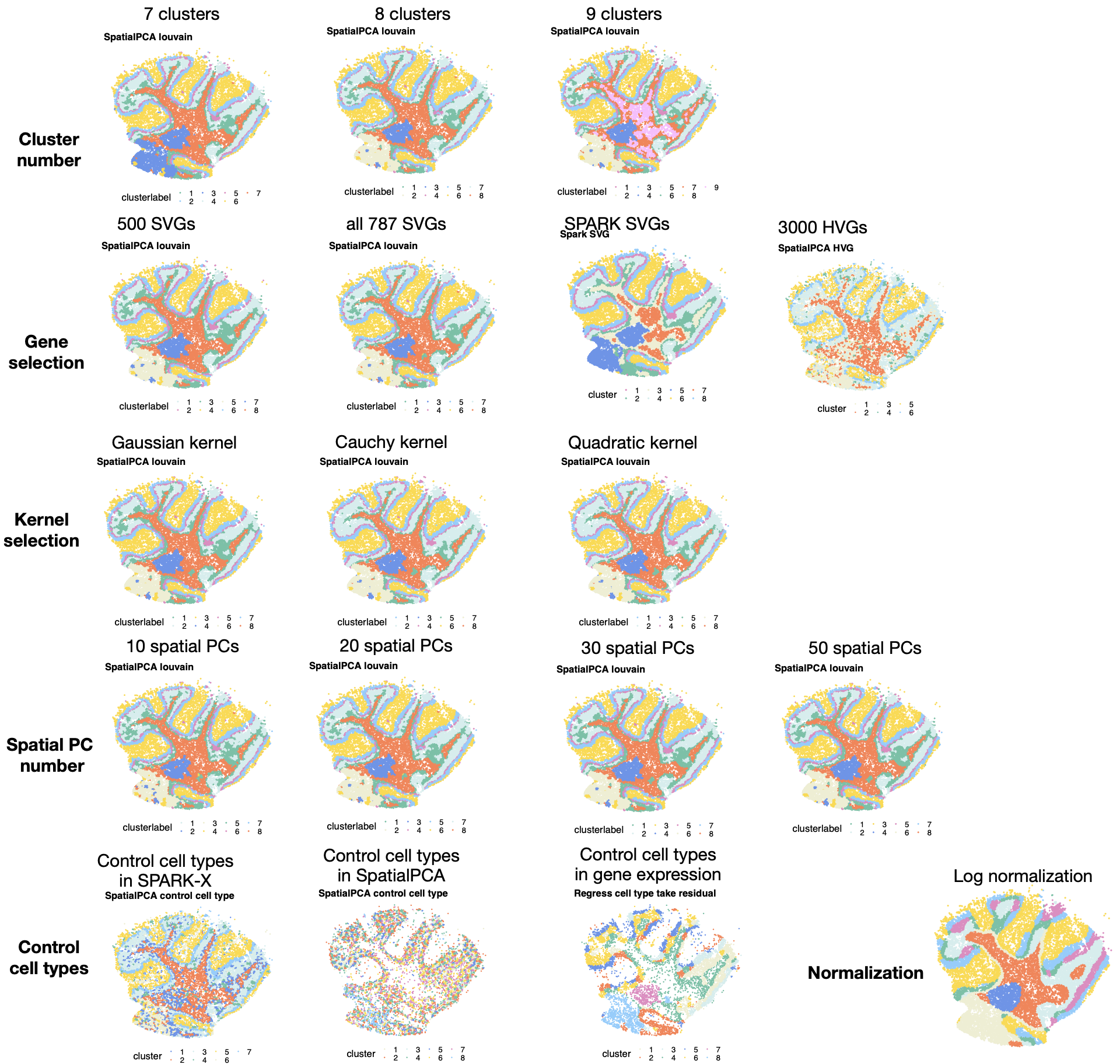

**Figure S10. Sensitivity analysis in the Slide-seq data**. **First row**: clustering results obtained with a different cluster number, which is set to be either 7, 8, and 9. Increasing number of clusters leads to more refined tissue structures. **Second row**: clustering results obtained using a different set of input genes, include top 500 SVGs, all SVGs detected by SPARK-X, all SVGs detected by SPARK, and top 3000 HVGs. In SPARK and SPARK-X, the adjusted p value cut-off for determining the SVGs is set to be 0.05. **Third row:** clustering results based on spatial PCs extracted using either the Gaussian kernel, the Cauchy kernel, or the rational quadric kernel. **Fourth row**: clustering results obtained using either the top 10, 20, 30, or 50 spatial PCs. **Fifth row**: left: clustering results obtained by controlling cell types when selecting SVGs in SPARK-X; controlling cell types in SpatialPCA; controlling cell types by regressing them out from the input gene expression and take the residuals. Right: clustering results obtained by SpatialPCA with gene expression normalized through log normalization.

Figure S11

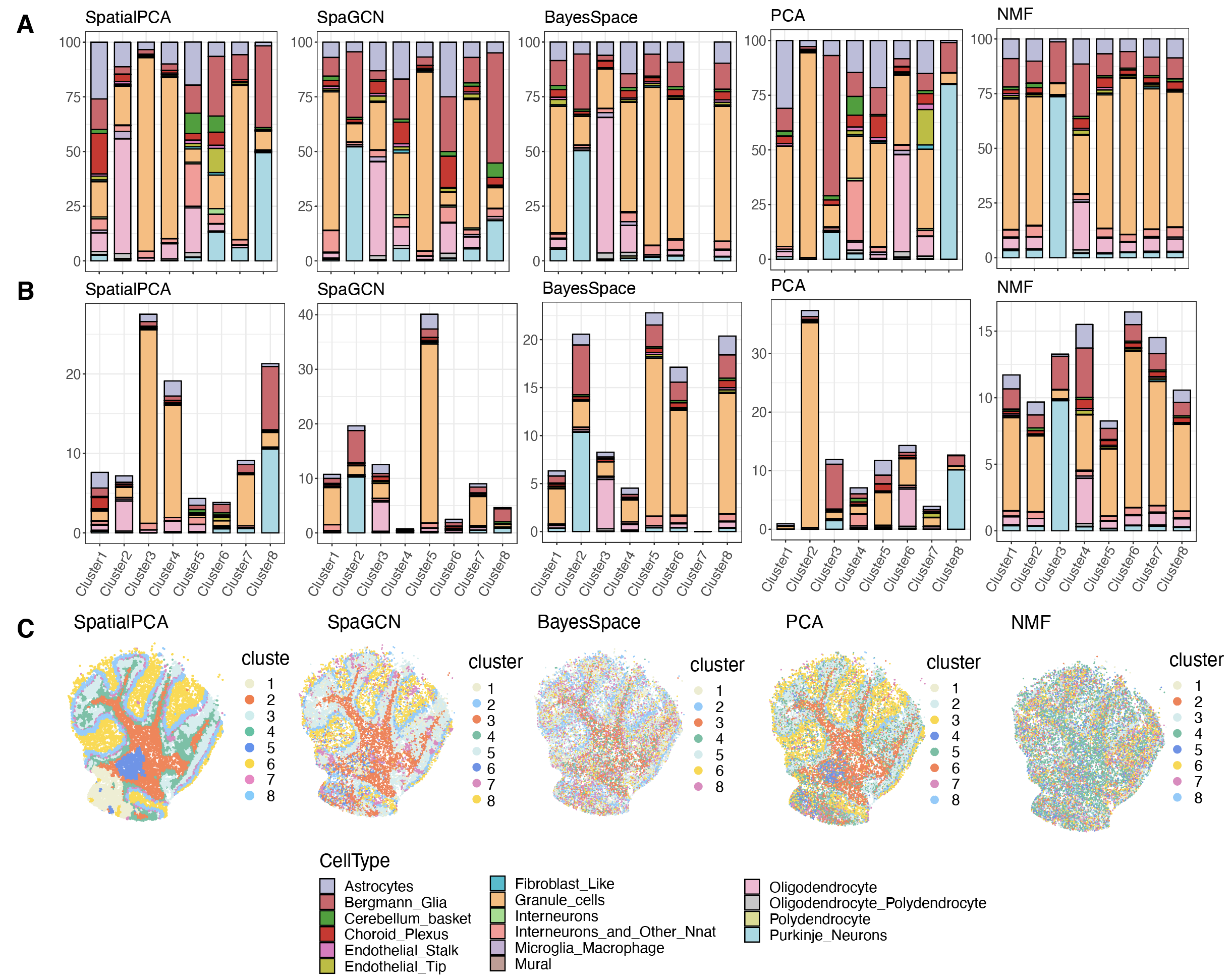

**Figure S11.** **Comparison of the cell type composition of the spatial domains detected by different methods in the Slide-seq data**. The percentage of cell types annotated (y-axis) is shown on each tissue domain (x-axis) detected by different methods. Examined methods include SpatialPCA, SpaGCN, BayesSpace, PCA, and NMF. (**A**): results are scaled with respect to each spatial domain, such that the summation of the cell type percentages in each domain is 100%. (**B**): results are scaled with respect to the cell types, such that the summation of all cell types across all tissue regions is 100%. (**C**): Clustering results in each method. The clustering labels correspond to the x-axis in the left and middle panel.

Figure S12

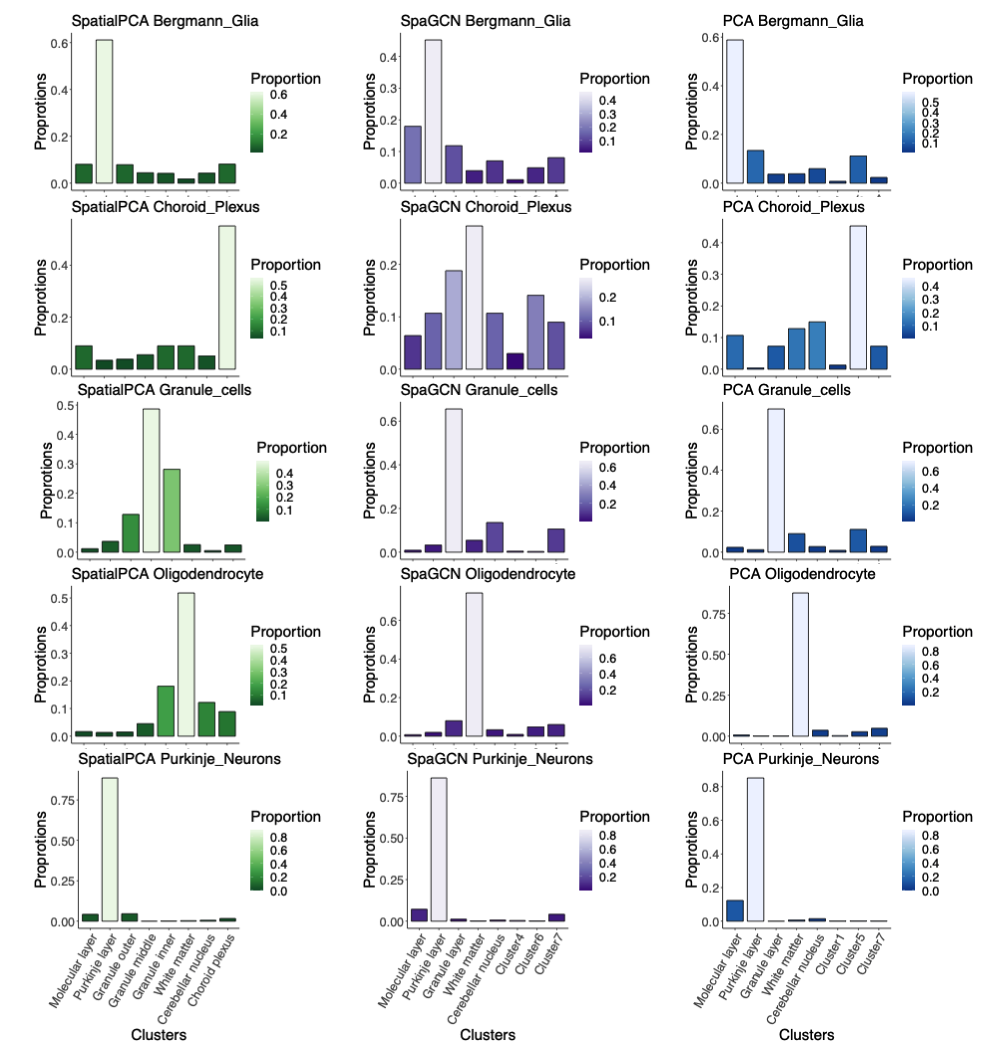

**Figure S12. Distribution of cell types in each cluster for different methods in the Slide-seq data.** The summation of the cell type percentages in all clusters is 100% for SpatialPCA (left panel), SpaGCN (middle panel), and PCA (right panel). The Bergmann glia cells and Purkinje neurons are located in the Purkinje layer, the choroid plexus cells are located in the choroid plexus, the granule cells are located in the granule cell layer, the oligodendrocyte cells are located in white matter.

Figure S13

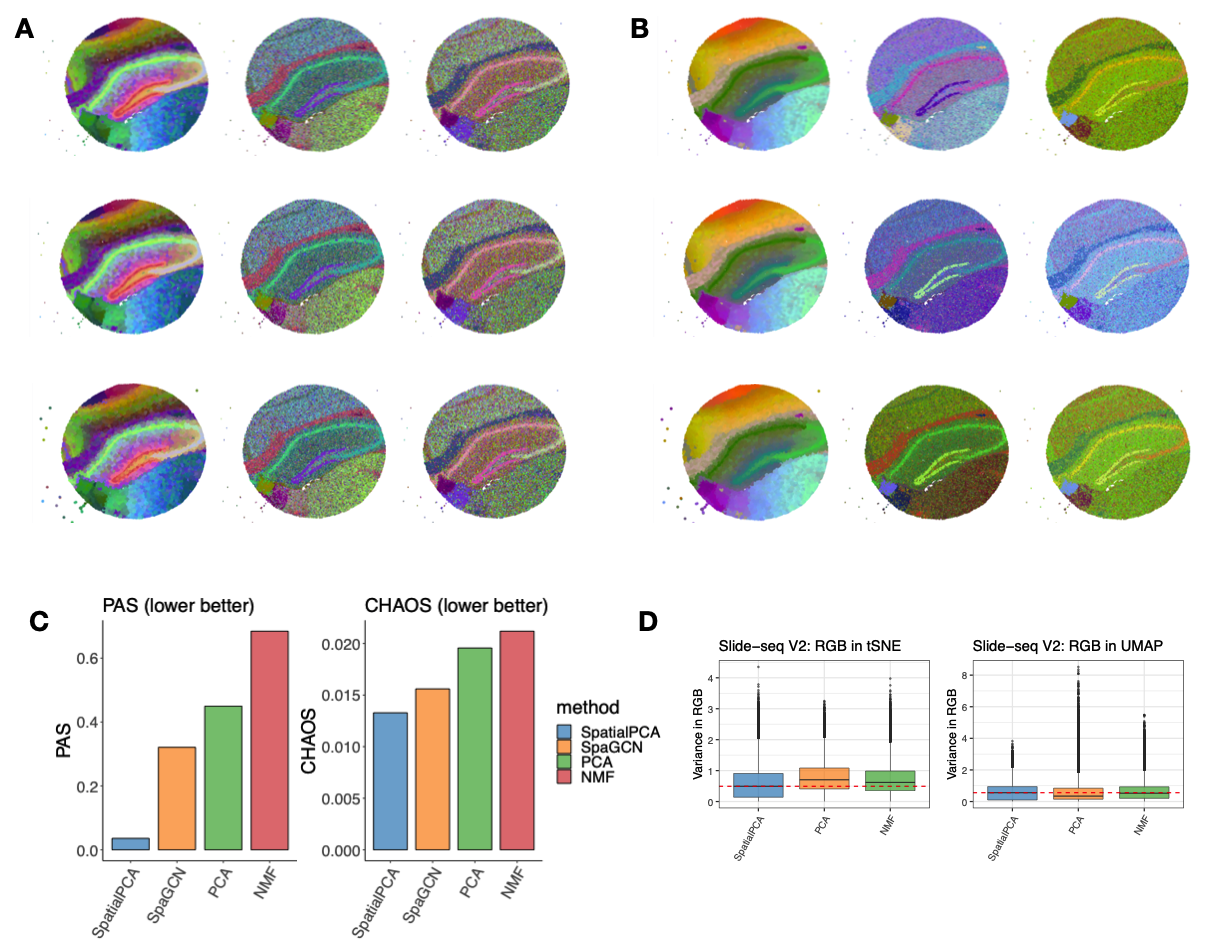

**Figure S13.** **RGB plots for the Slide-seq V2 data**. For SpatialPCA, PCA, and NMF, we summarized the inferred low dimensional components into three UMAP (**A**) or tSNE components (**B**) and visualized the three resulting components with red/green/blue (RGB) colors through the RGB plot. The RGB plot from SpatialPCA displays tissue structure organization of the hippocampus region and show less color differences within a local area. We also scaled up spatial PCs/regular PCs 10 times (second row) and 20 times (third row) to see the influence of range of the PCs to RGB plots, the tSNE/UMAP results and RGB plots have very similar patterns as shown at the original scale (first row). (**C**) The PAS and CHAOS scores (the lower the better) of clustering results in SpatialPCA is the lowest compared with BayesSpace, SpaGCN, PCA and NMF. (**D**) The weighted RGB values in SpatialPCA have lower variance than PCA or NMF in nearby spots.

Figure S14

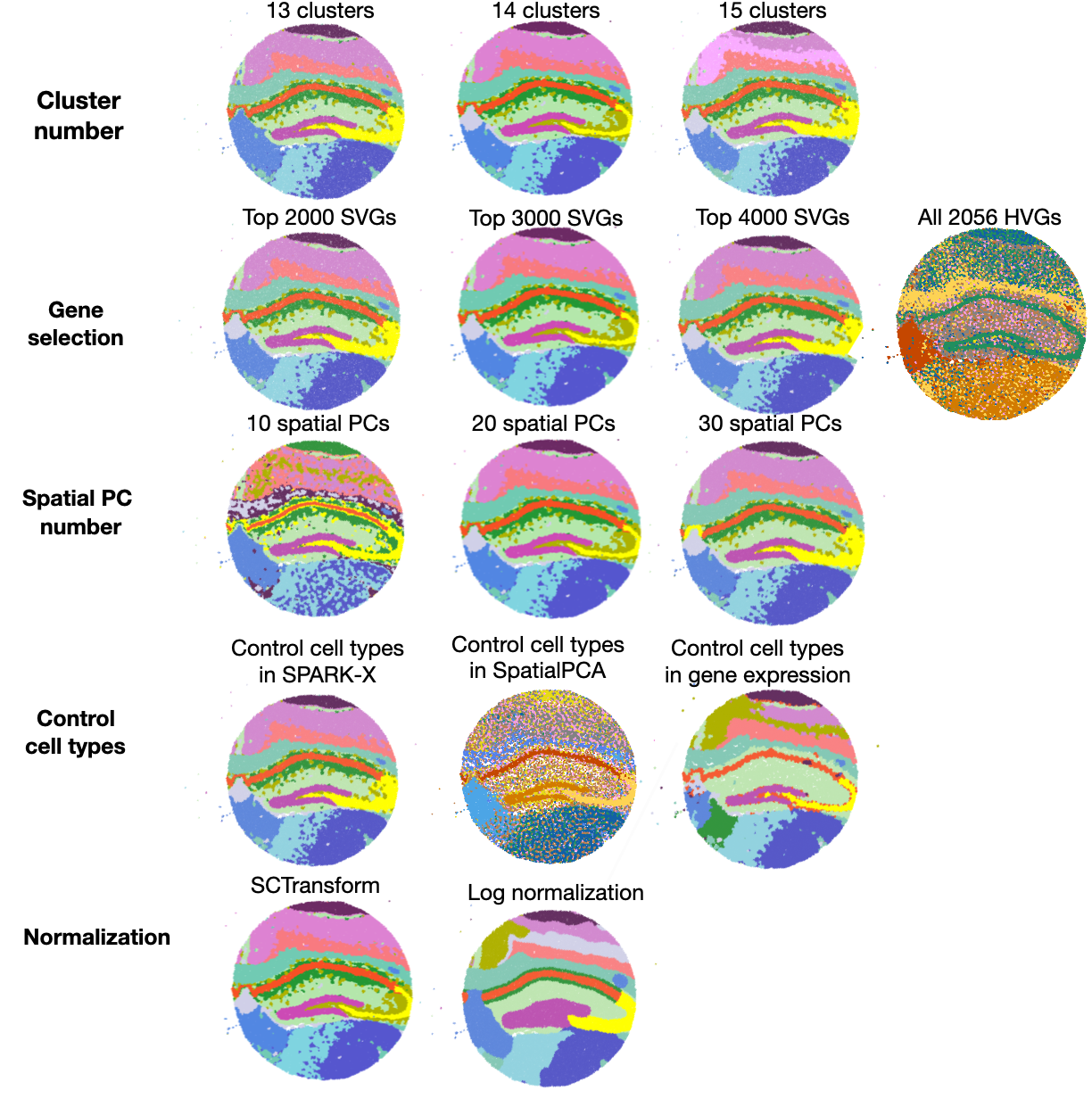

**Figure S14. Sensitivity analysis in the Slide-seq V2 data**. **First row**: clustering results obtained with a different cluster number, which is set to be either 13, 14, and 15. Increasing number of clusters leads to more refined tissue structures. **Second row**: clustering results obtained using a different set of input genes, include top 2000 SVGs, top 3000 SVGs, and top 4000 SVGs detected by SPARK-X, and all 2056 HVGs. In SPARK-X, the adjusted p value cut-off for determining the SVGs is set to be 0.05. **Third row:** clustering results obtained using either the top 10, 20, or 30 spatial PCs. **Fourth row**: clustering results obtained by controlling cell types when selecting SVGs in SPARK-X; controlling cell types in SpatialPCA; controlling cell types by regressing them out from the input gene expression and take the residuals. **Fifth row**: clustering results obtained by SpatialPCA with gene expression normalized through SCTransform normalization or log normalization.

Figure S15

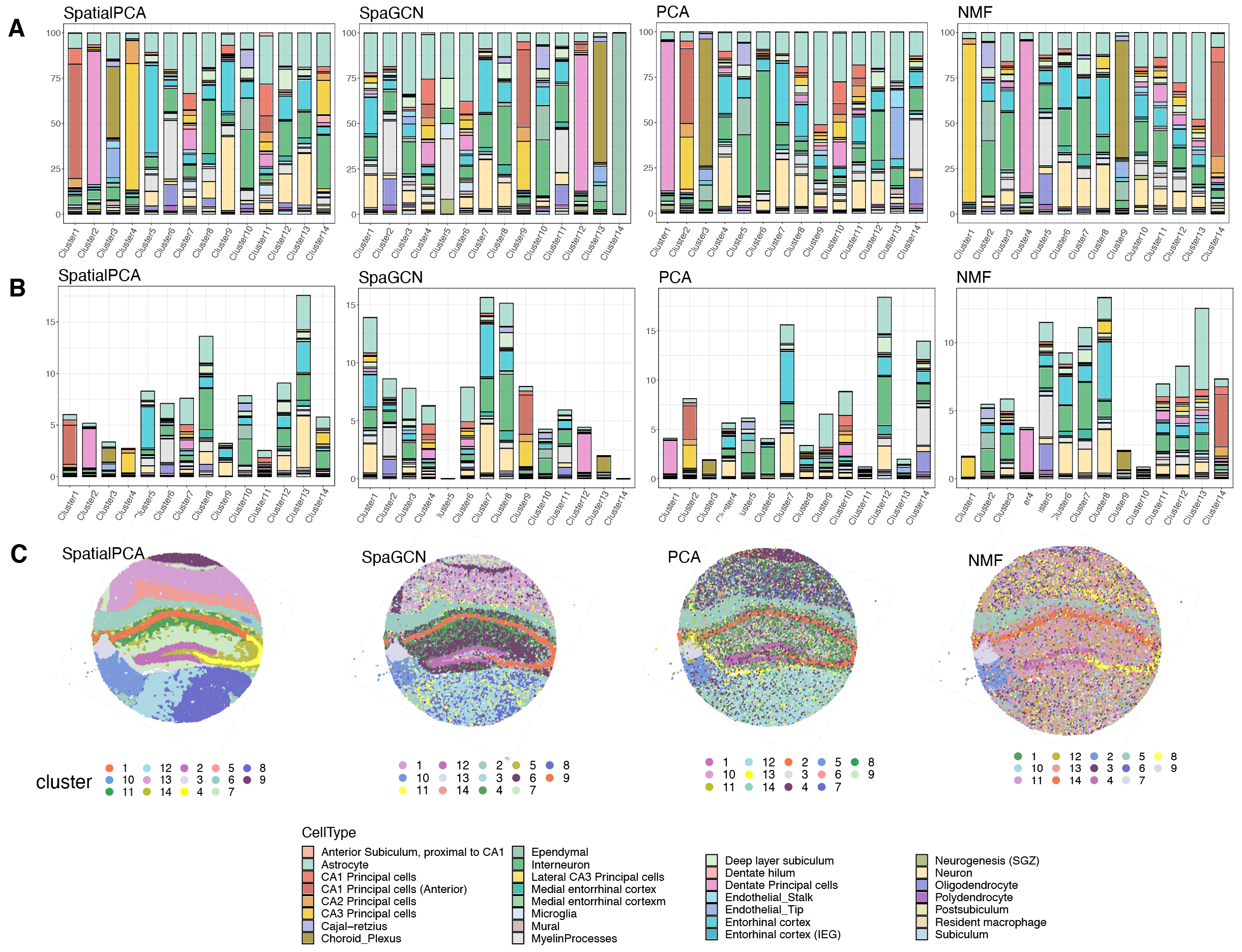

**Figure S15.** **Comparison of the cell type composition of the spatial domains detected by different methods in the Slide-seq V2 data**. The percentage of cell types annotated (y-axis) is shown on each tissue domain (x-axis) detected by different methods. Examined methods include SpatialPCA, SpaGCN, PCA, and NMF. (**A**): results are scaled with respect to each spatial domain, such that the summation of the cell type percentages in each domain is 100%. (**B**): results are scaled with respect to the cell types, such that the summation of all cell types across all tissue regions is 100%. (**C**): Clustering results in each method. The clustering labels correspond to the x-axis in (B).

Figure S16

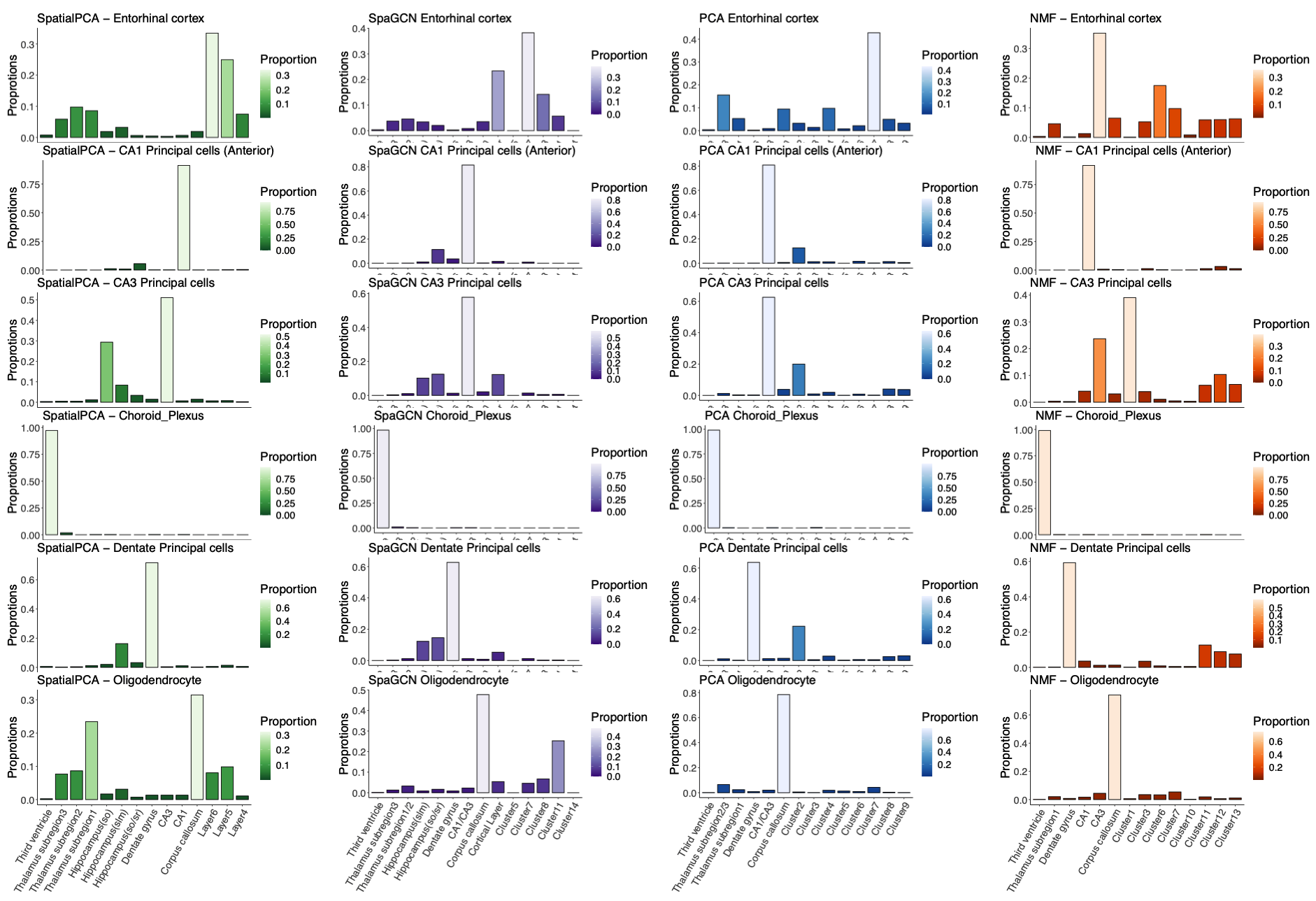

**Figure S16. Distribution of cell types in each cluster for different methods in Slide-seq V2 data.** The summation of the cell type percentages in all clusters is 100% for SpatialPCA (first column), SpaGCN (second column), PCA (third column) and NMF (fourth column). The entorhinal cortex cells are located in the cortical layers 4-6; the CA1 principal cells (anterior) are located in the CA1 region; the CA3 principal cells are located in the CA3 region; the choroid plexus are located in the third ventricle; the dentate principle cells are located in the dentate gyrus; and the oligodendrocyte are located in the corpus callosum as detected by SpatialPCA.

Figure S17

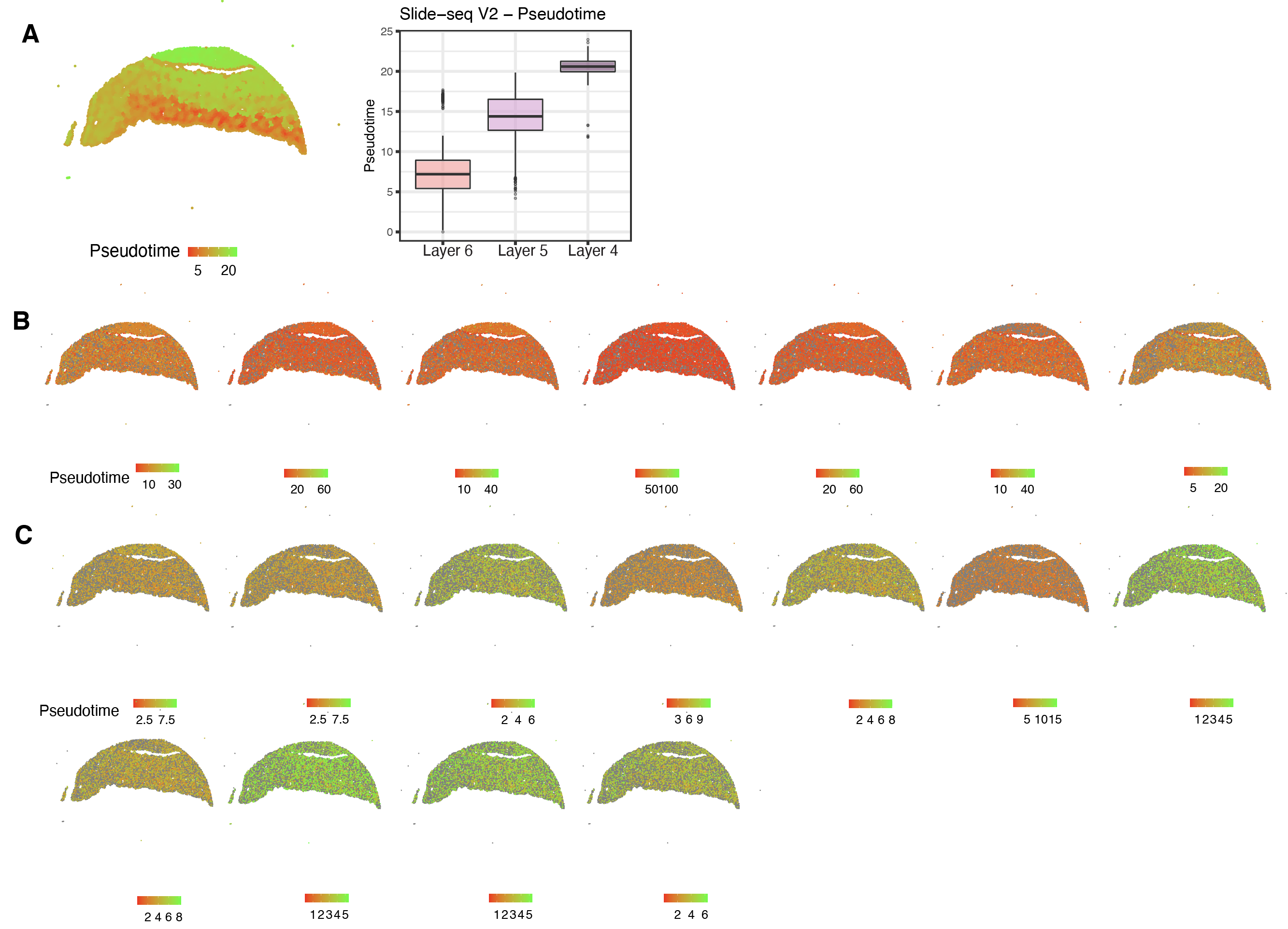

**Figure S17.** **Spatial trajectory inference results in the cortical layers of the Slide-seq V2 data**. (**A**) Left: Visualizaton of the pseudo-time for the inferred trajectory in cortical layers of the Slide-seq V2 data in SpatialPCA. Right: Boxplot showing the pseudotime of locations in cortical layer 4, 5, and 6. (**B**) Visualizaton of the inferred pseudo-time for seven trajectories in cortical layers of the Slide-seq V2 data in PCA. (**C**) Visualizaton of the inferred pseudo-time for eleven trajectories in cortical layers of the Slide-seq V2 data in NMF.

Figure S18

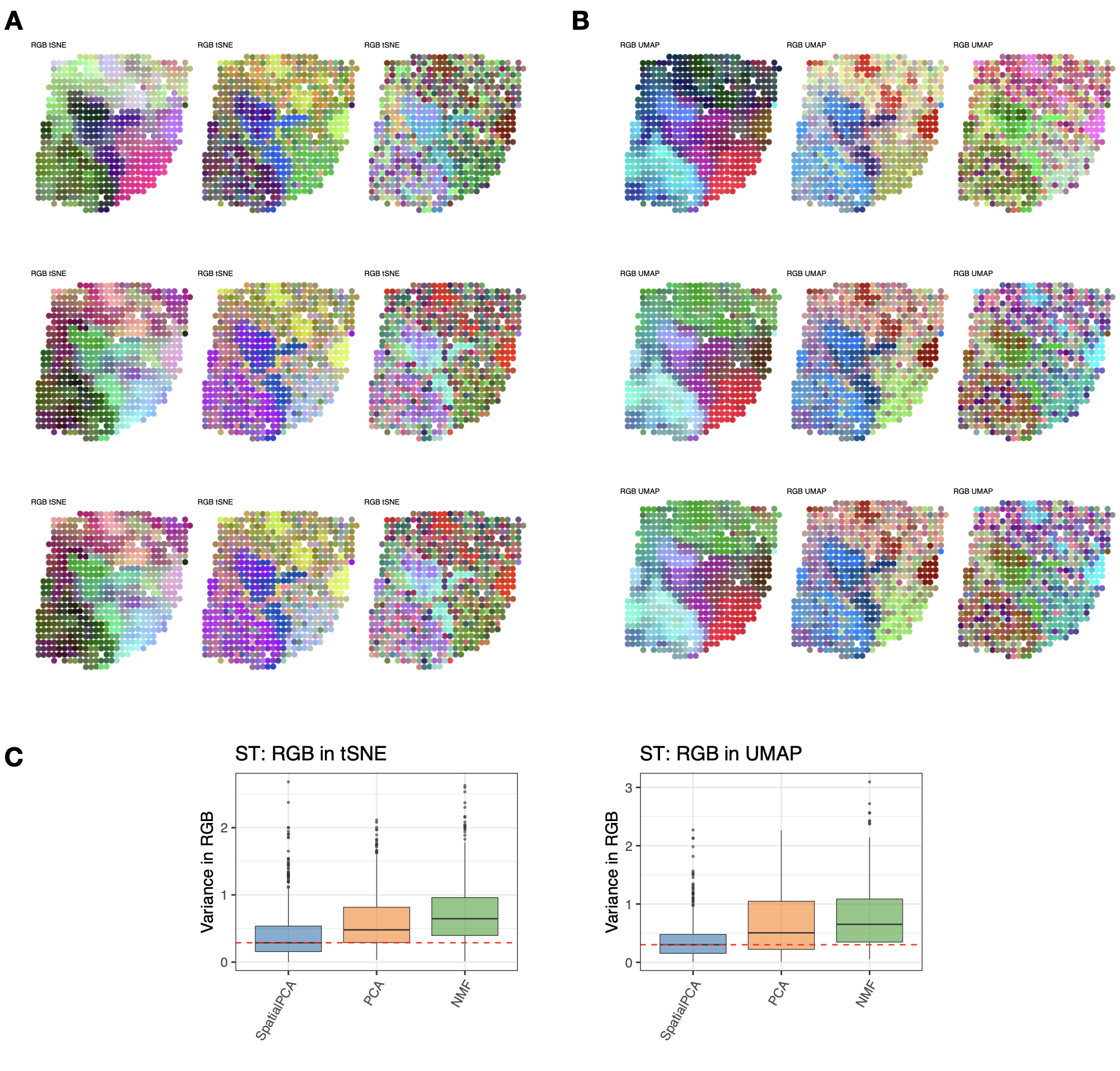

**Figure S18.** **RGB plots for the ST data**. For SpatialPCA, PCA, and NMF, we summarized the inferred low dimensional components into three tSNE components (**A**) and UMAP (**B**) and visualized the three resulting components with red/green/blue (RGB) colors through the RGB plot. The RGB plot from SpatialPCA displays tissue structure organization of the breast tumor and show less color differences within a local area. We also scaled up spatial PCs/regular PCs 10 times (second row) and 20 times (third row) to see the influence of range of the PCs to RGB plots, the tSNE/UMAP results and RGB plots have very similar patterns as shown at the original scale (first row). (**C**) The weighted RGB values in SpatialPCA have lower variance than PCA or NMF in nearby spots.

Figure S19

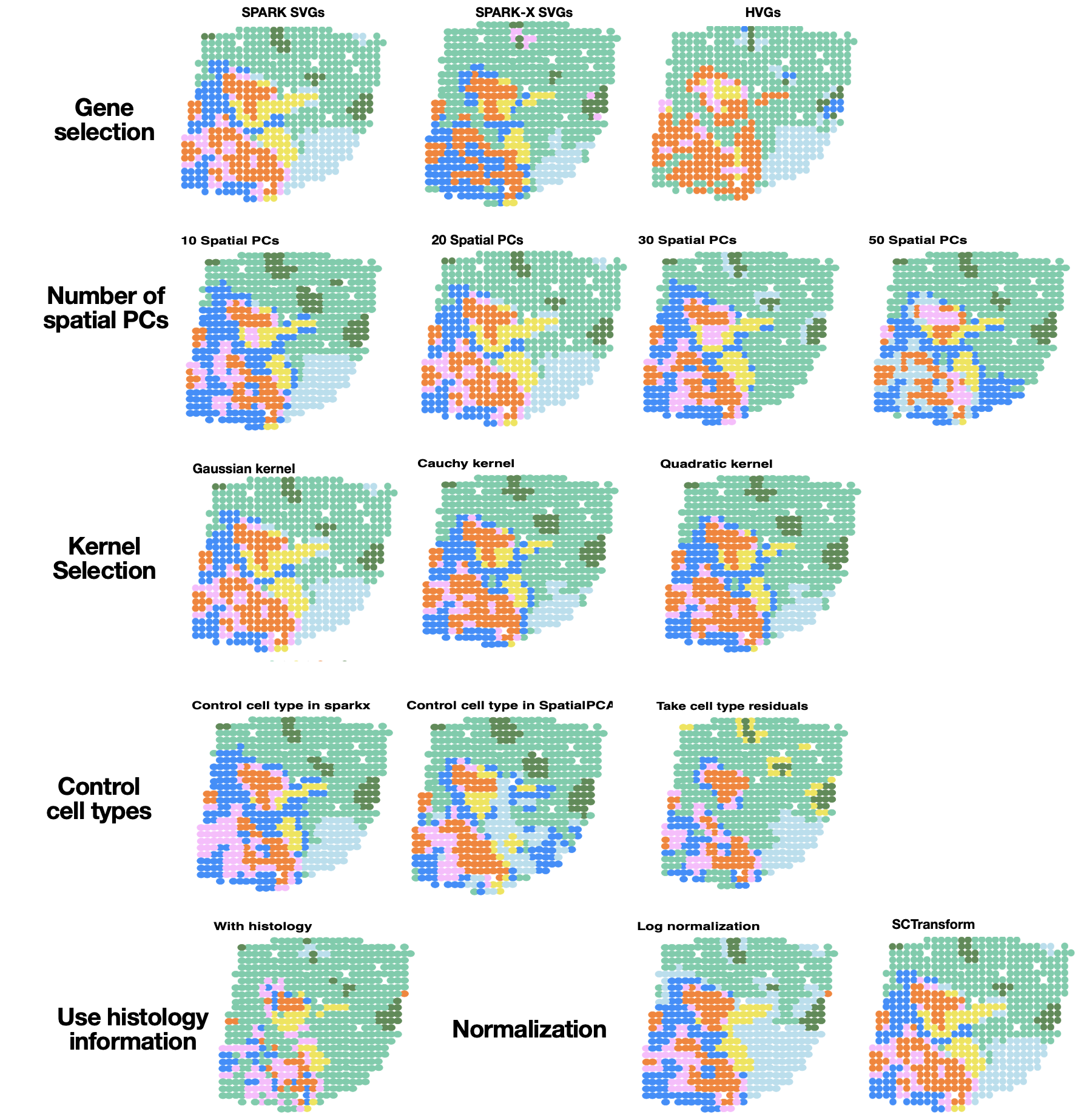

**Figure S19. Sensitivity analysis in the ST data**. **First row**: clustering results obtained using a different set of input genes, include SVGs detected by SPARK, SVGs detected by SPARK-X, and HVGs. In SPARK and SPARK-X, the adjusted p value cut-off for determining the SVGs is set to be 0.05. **Second row**: clustering results obtained using either the top 10, 20, 30, or 50 spatial PCs. **Third row**: clustering results based on spatial PCs extracted using either the Gaussian kernel, the Cauchy kernel, or the rational quadric kernel. **Fourth row**: clustering results obtained by controlling cell types when selecting SVGs in SPARK-X; controlling cell types in SpatialPCA; controlling cell types by regressing them out from the input gene expression and take the residuals. **Fifth row**: Left: clustering results obtained by taking the histology information as a third dimension in location matrix. The histology information was extracted from the H&E image following SpaGCN. Right: clustering results obtained by SpatialPCA with gene expression normalized through SCTransform normalization or log normalization.

Figure S20

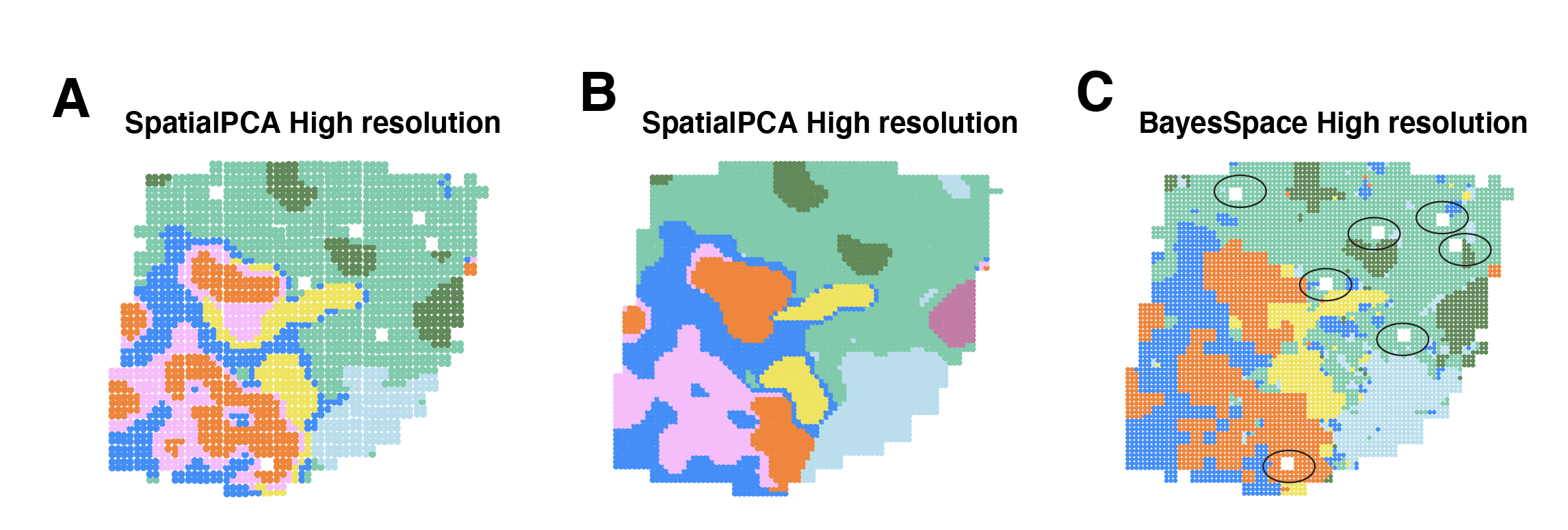

**Figure S20.** Visualization of high-resolution spatial map prediction in SpatialPCA and BayesSpace. (**A**) We generate 2,428 new locations by adding a small distance to the x and y coordinates of the original location to obtain four new locations. We performed clustering on the predicted spatial PCs on the new locations. (**B**) We generate 4,933 new locations that cover the whole tissue section. We performed clustering on the predicted spatial PCs on the new locations. (C) Visualization of high-resolution spatial map prediction in BayesSpace by its default setting. BayesSpace generates high resolution prediction map based on the existing spots, when there are spots filtered out due to low expression counts in the original data, its reconstructed high resolution map would also have gaps around those spots. We circled the unimputed spots in BayesSpace.

Figure S21

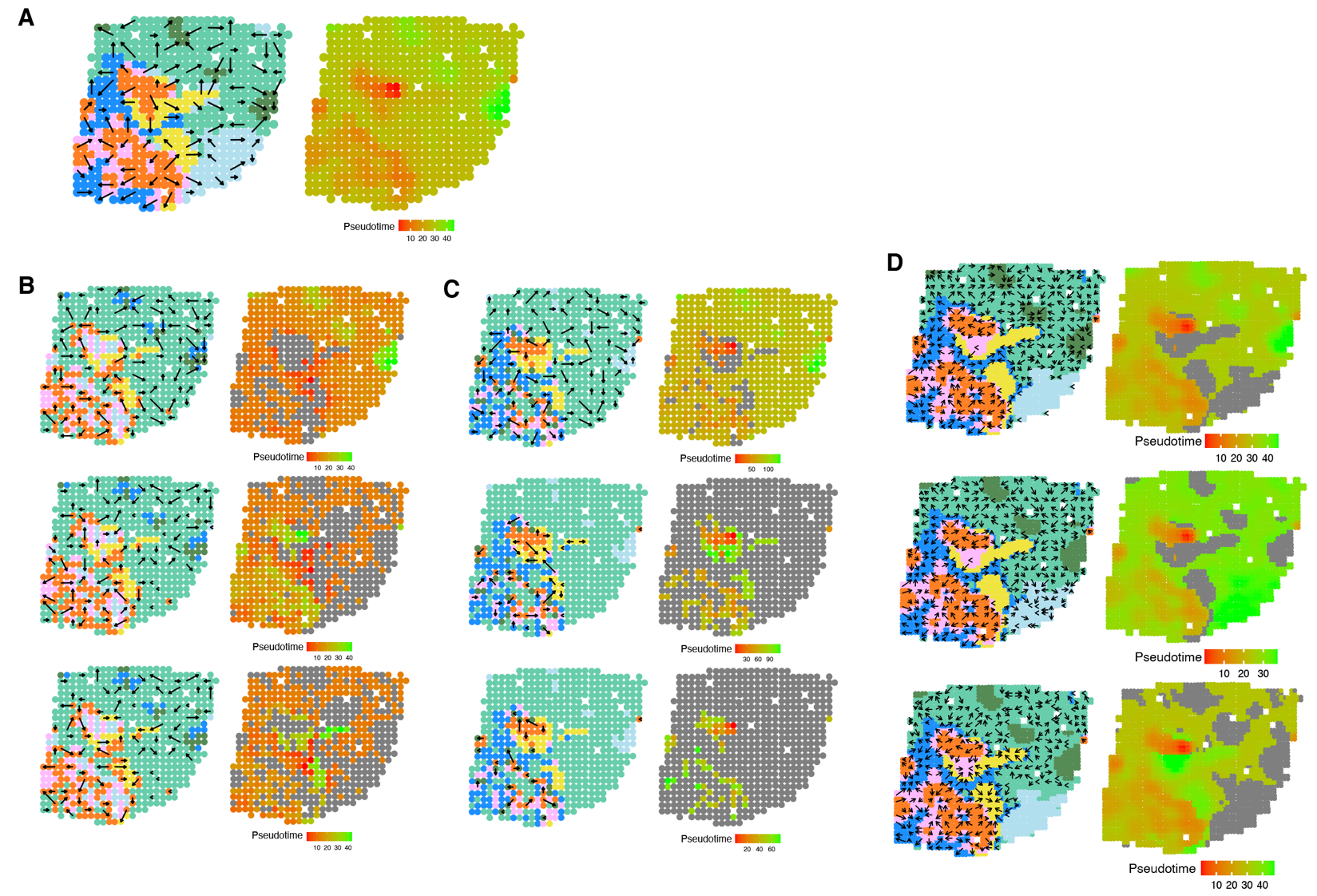

**Figure S21. Spatial trajectory inference results in the ST tumor data**. **(A)**: Visualization of the trajectory inferred by SpatialPCA in the original data. **(B)**: Visualization of the trajectory inferred by PCA. (**C**) Visualization of the trajectory inferred by NMF. (**D**): Visualizaton of the three trajectories inferred by SpatialPCA on the SpatialPCA constructed high-resolution spatial map. In all panels, arrows point from tissue locations with low pseudo-time to tissue locations with high pseudo-time. Color represents different tissue regions.

Figure S22

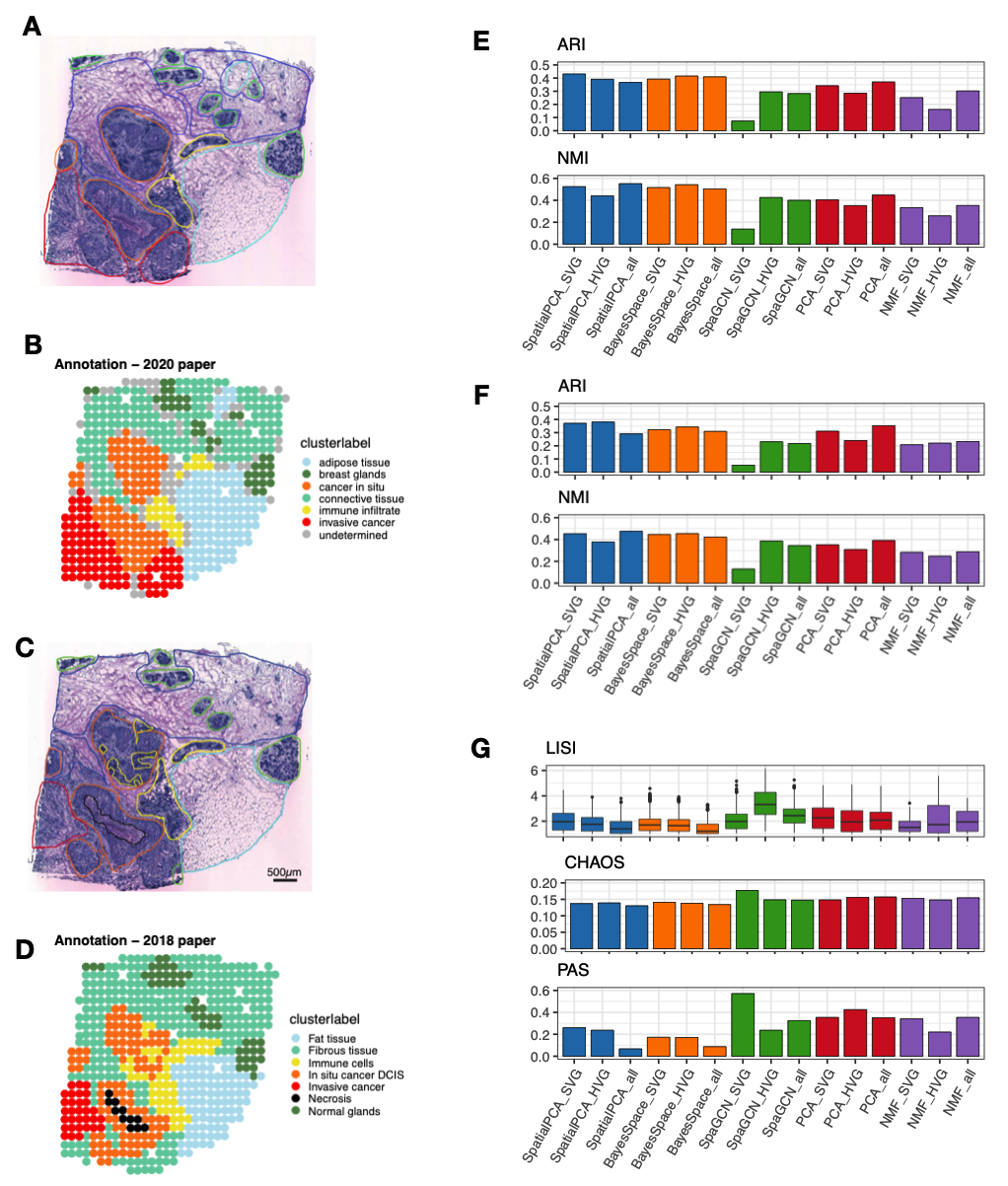

**Figure S22.** **Clustering results obtained based on different methods in the ST data.** (**A**) Haematoxylin and eosin (H&E) staining image shows distinct tissue regions annotated by a pathologist in the original study (Andersson et al. 2021). The annotated tissue regions include invasive cancer (red), adipose tissue (cyan), connective tissue (light green), breast glands (green), cancer in situ (orange), and immune infiltrate (yellow). (**B**) Visualization of pathologist hand annotation in the original study (Andersson et al. 2020). (**C**) Haematoxylin and eosin (H&E) staining image shows distinct tissue regions annotated by a pathologist in a previous version of the original study (Salmén et al. 2018). The annotated tissue regions include invasive cancer (red), fat tissue (cyan), fibrous tissue (blue), normal breast glands (green), in situ cancer/DCIS (orange), immune cells (yellow), and necrosis (black). (**D**) Visualization of hand annotation of spots based on the H&E annotation. (**E**) Clustering results measured by ARI (the higher the better) and NMI (the higher the better) in the original study (Andersson et al. 2021). (**F**) Clustering results measured by ARI (the higher the better) and NMI (the higher the better) in the previous study (Salmén et al. 2018). (**G**) Clustering results measured by LISI (the lower the better), PAS (the lower the better), and CHAOS (the lower the better). In dimension reduction methods (SpatialPCA, PCA, and NMF), clustering was performed based on the inferred low-dimensional components. For spatial domain clustering methods (BayesSpace and SpaGCN), clustering was performed based the default settings. All the methods are paired with SVGs, HVGs and all genes.

Figure S23

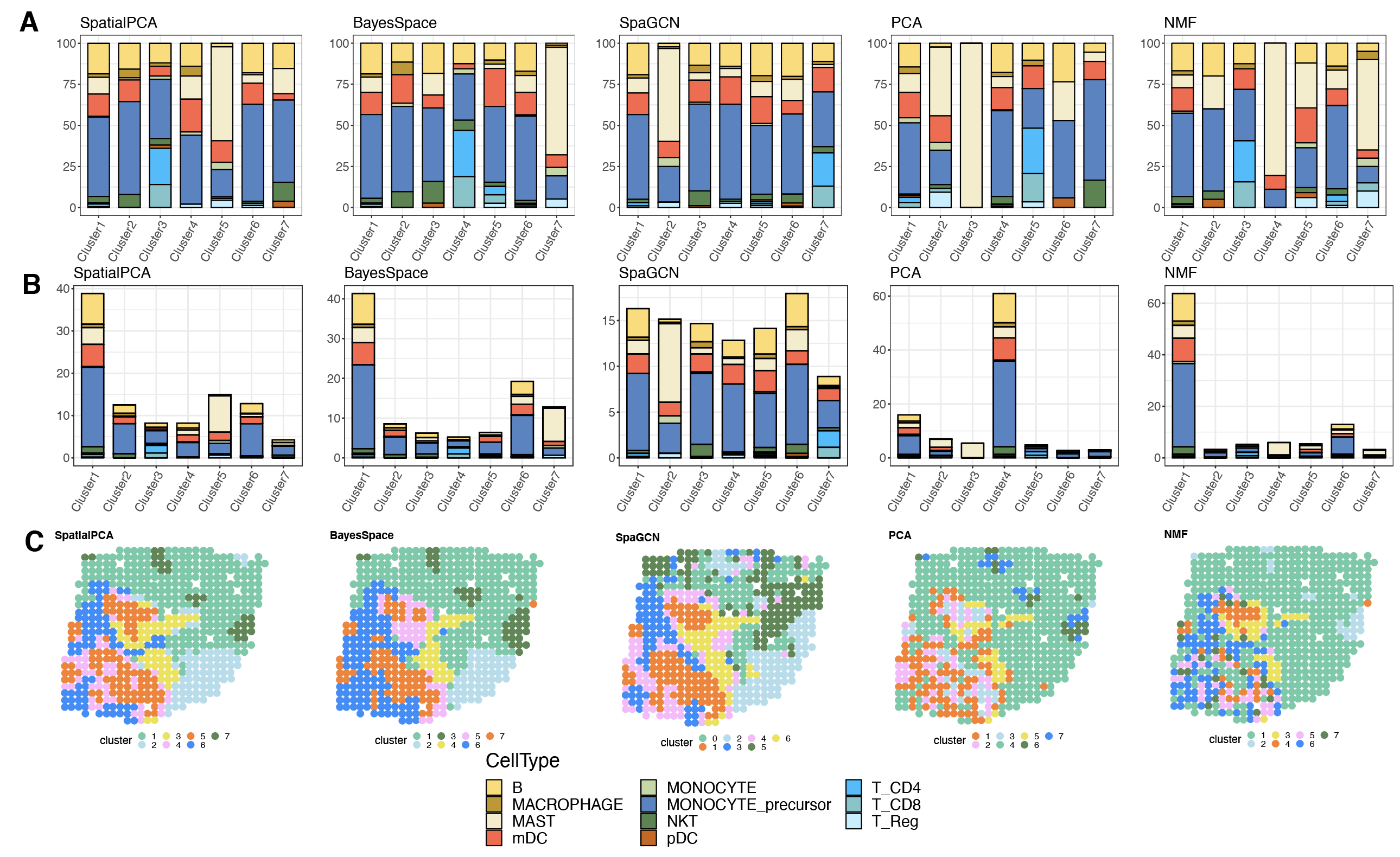

**Figure S23.** **Comparison of the cell type composition of the spatial domains detected by different methods in the ST tumor data**. The percentage of cell types annotated (y-axis) is shown on each tissue domain (x-axis) detected by different methods. Examined methods include SpatialPCA, BayesSpace, SpaGCN, PCA, and NMF. (**A**): results are scaled with respect to each spatial domain, such that the summation of the cell type percentages in each domain is 100%. (**B**): results are scaled with respect to the cell types, such that the summation of all cell types across all tissue regions is 100%. (**C**): Clustering results in each method. The clustering labels correspond to the x-axis in the left and middle panel. The clustering labels correspond to the x-axis in the left and middle panel. The reference scRNA-seq data for ST data that contain multiple immune cell types is collected on breast cancer via the drop-seq based platform.

**Figure S24**

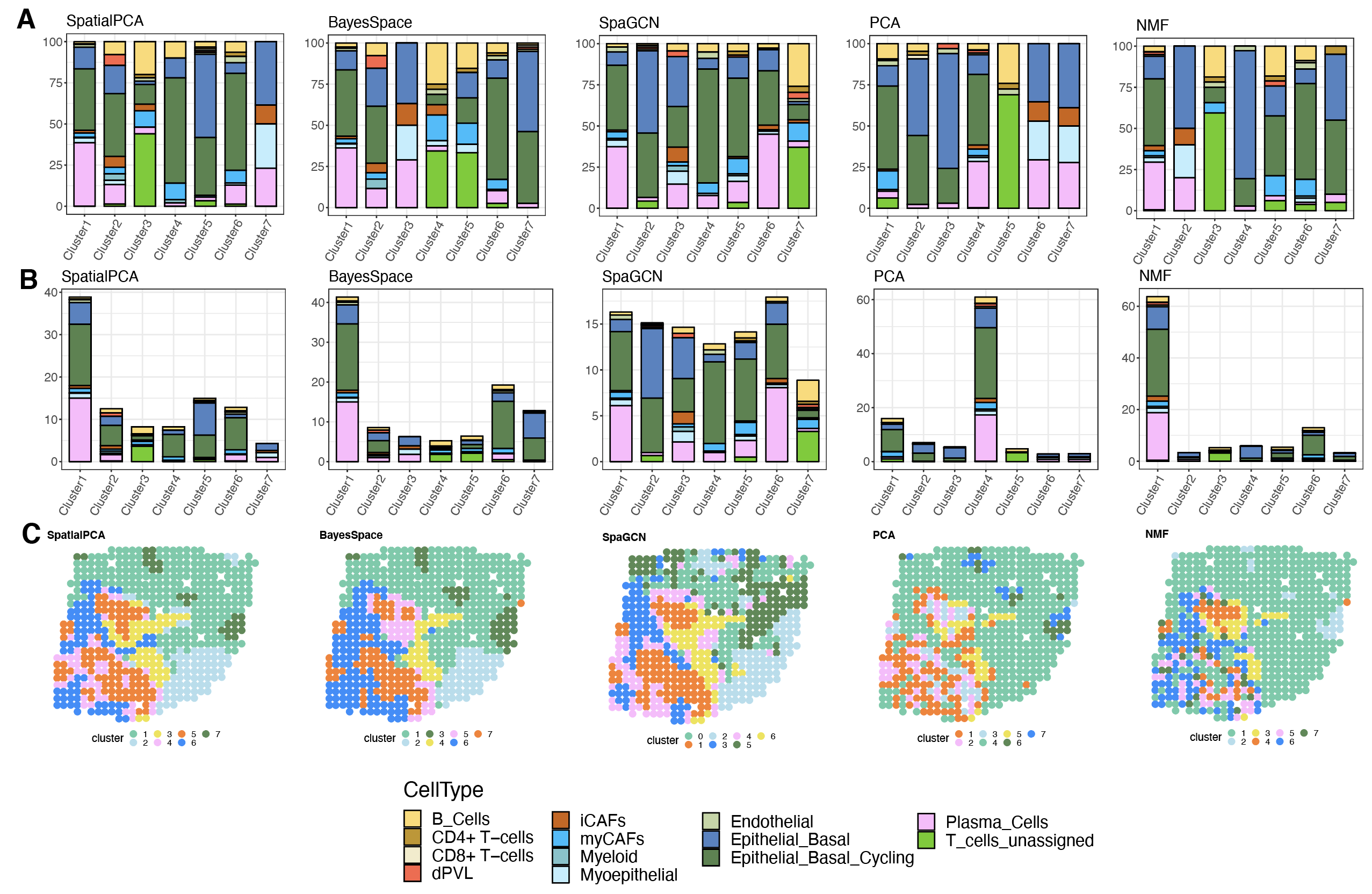

**Figure S24.** **Comparison of the cell type composition of the spatial domains detected by different methods in the ST tumor data**. The percentage of cell types annotated (y-axis) is shown on each tissue domain (x-axis) detected by different methods. Examined methods include SpatialPCA, BayesSpace, SpaGCN, PCA, and NMF. (**A**): results are scaled with respect to each spatial domain, such that the summation of the cell type percentages in each domain is 100%. (**B**): results are scaled with respect to the cell types, such that the summation of all cell types across all tissue regions is 100%. (**C**): Clustering results in each method. The clustering labels correspond to the x-axis in the left and middle panel. The clustering labels correspond to the x-axis in the left and middle panel. The reference scRNA-seq data for ST data that contain immune cell types and malignant cell types is collected on breast cancer via the InDrop platform.

Figure S25

DLPFC

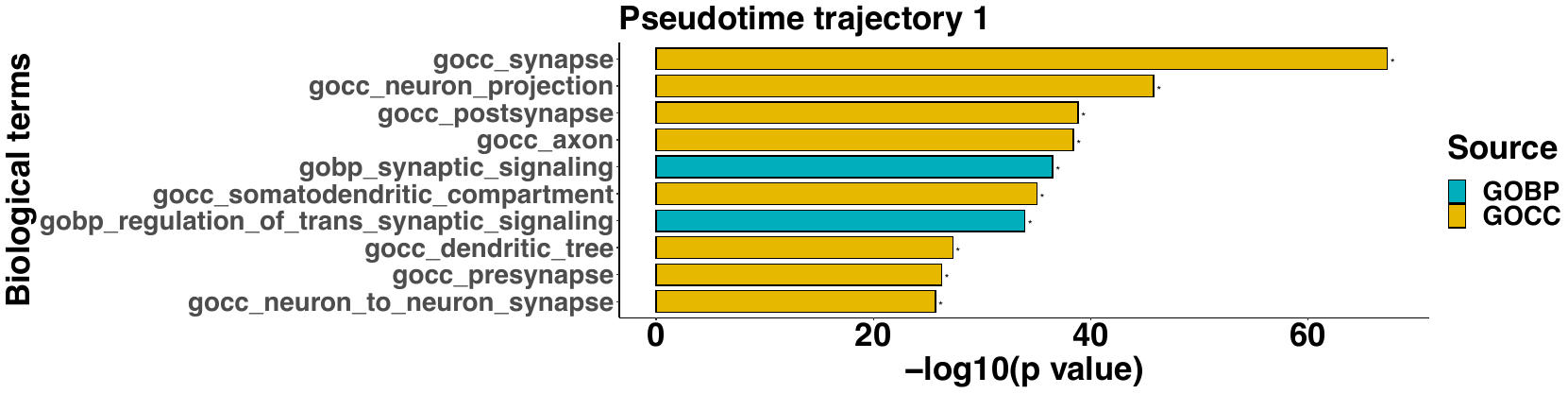

Slide-seq V2 cortical layers

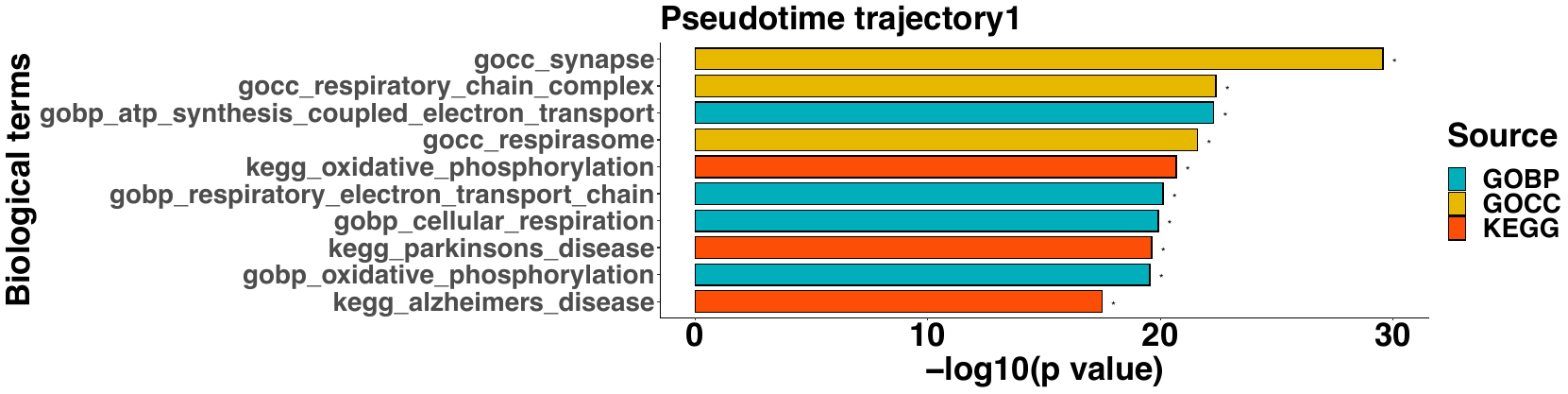

ST tumor

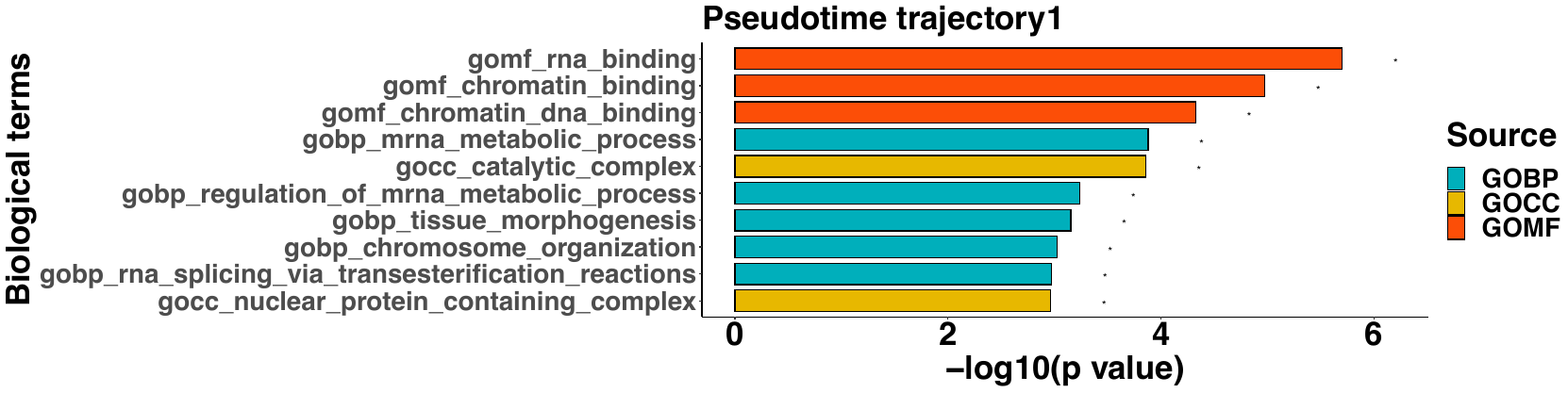

**Figure S25.** **Gene set** **enrichment analysis on the pseudo-time associated genes in the ST tumor data**. The top 10 enriched gene sets are shown for each of the three detected trajectories. Color represents different data sources for annotating the gene sets. (**A**) The enriched gene sets in DLPFC data. (**B**) The enriched gene sets in cortical layers of the Slide-seq V2 data. (**C**) The enriched gene sets in ST tumor data.

Figure S26

A.

| 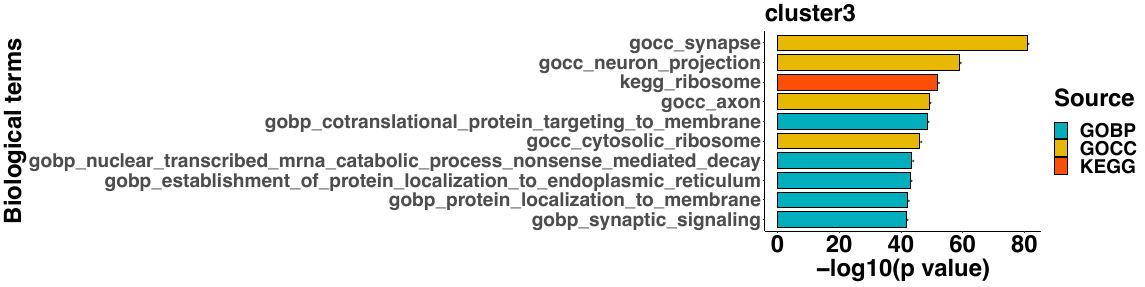 | 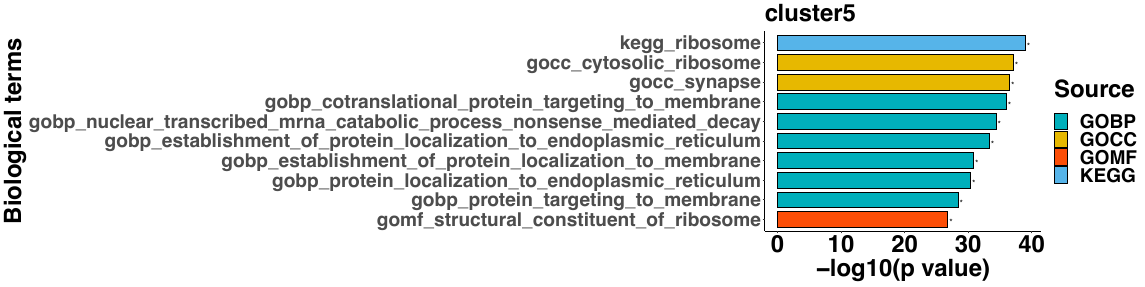 |
| --- | --- |
| 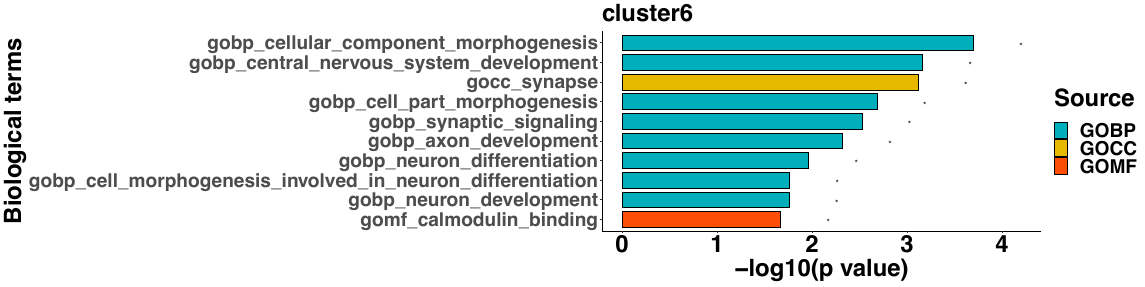 | 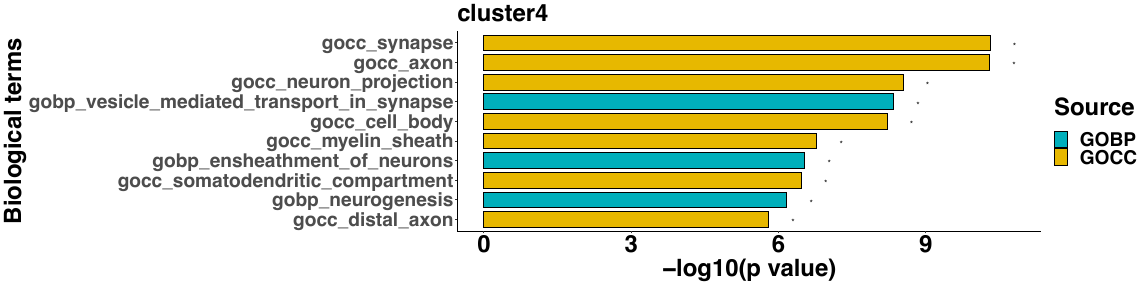 |
| 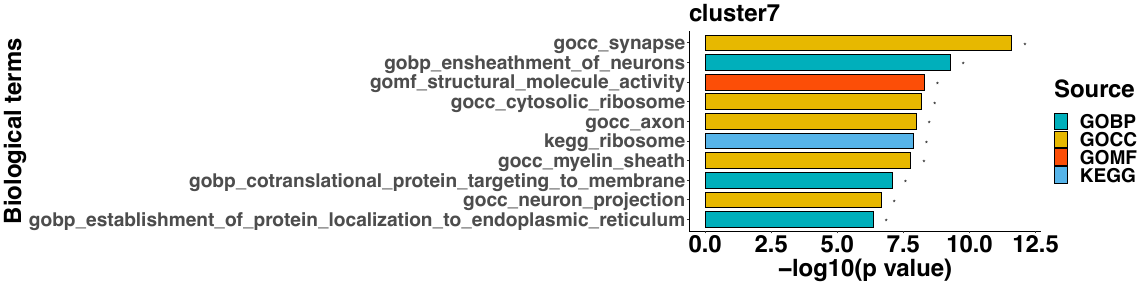 | 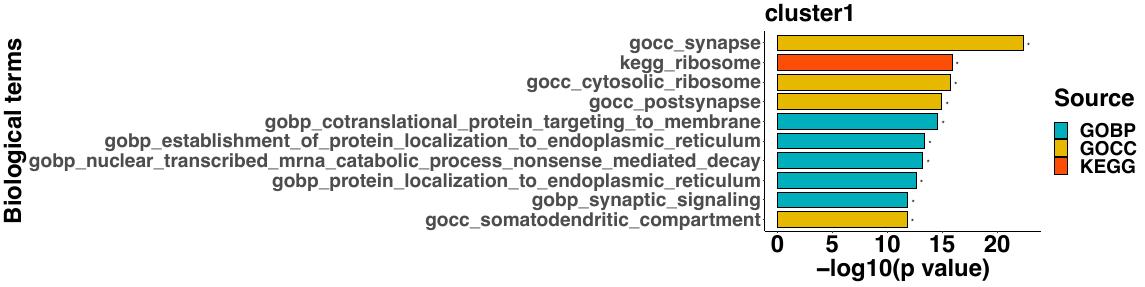 |
| 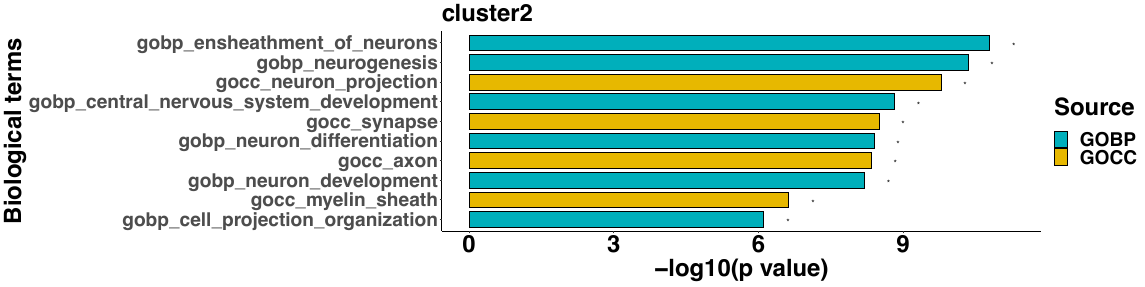 | 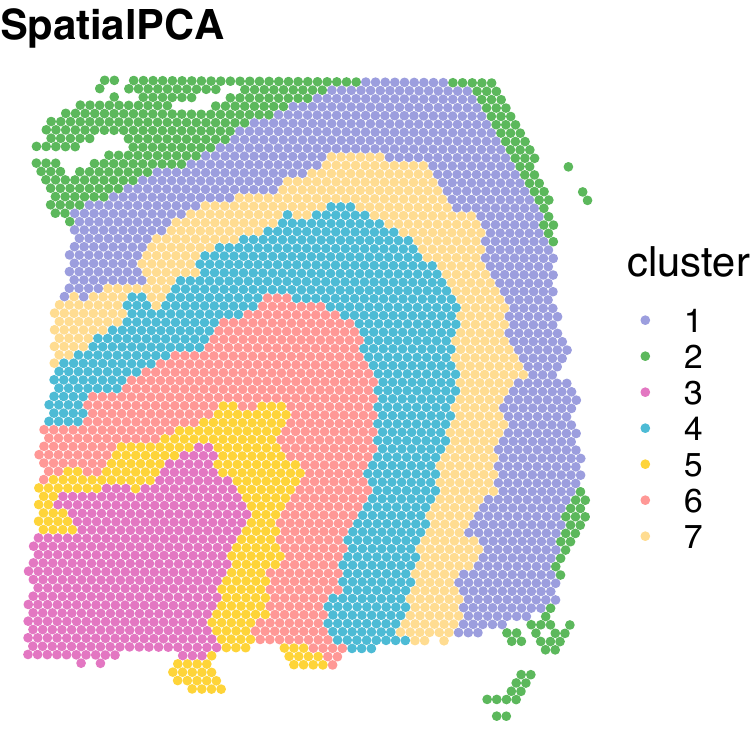 |

B.

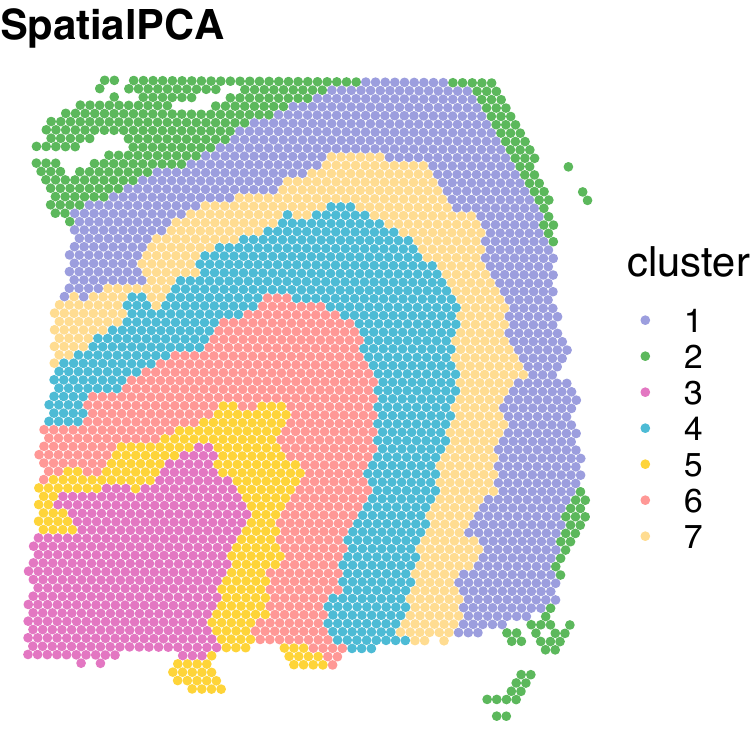

**Figure S26.** **Gene set enrichment analysis on the region-specific genes in the DLPFC data**. (**A**) The top 10 enriched gene sets are shown for each of the eight detected tissue regions. Color represents different data sources for annotating the gene sets. The cluster annotations are ordered from the inner most layer to outer most layer as shown in (**B**).

Figure S27

| 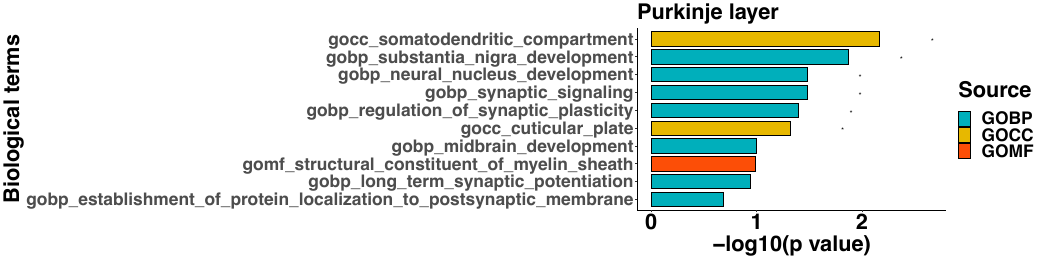 | 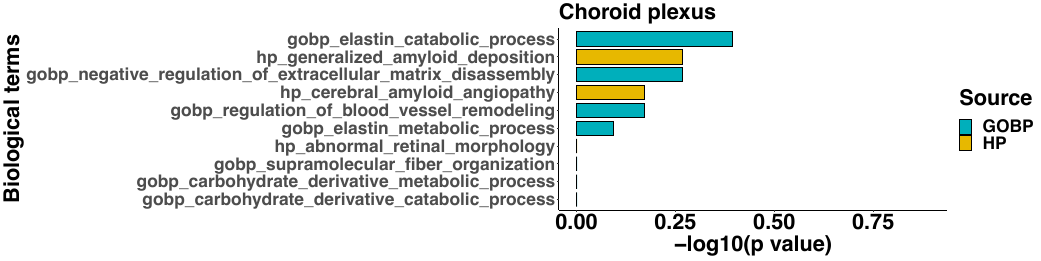 |
| --- | --- |
| 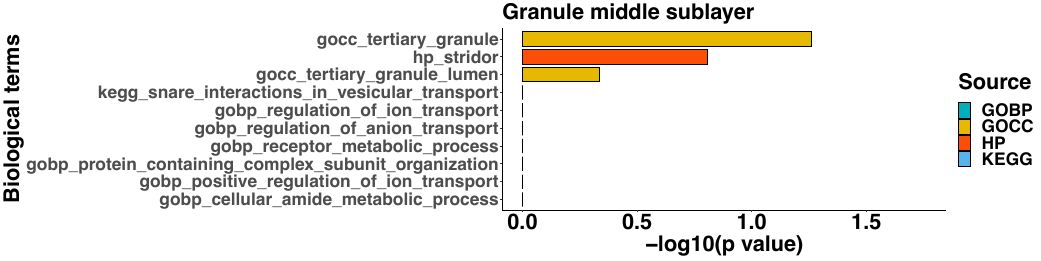 | 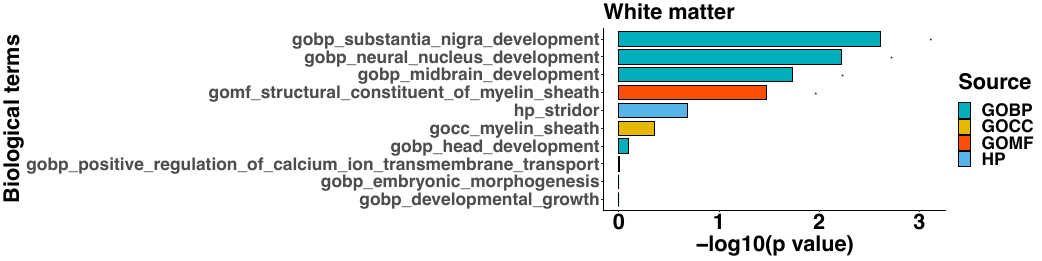 |
| 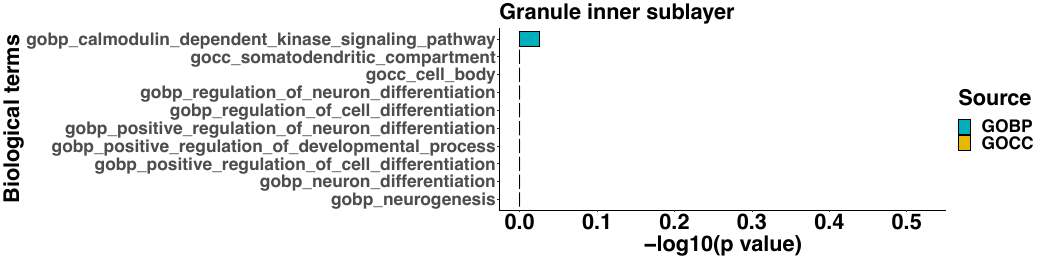 | 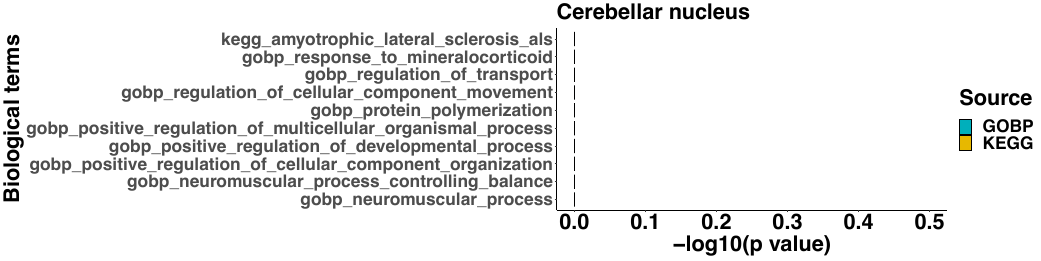 |
| 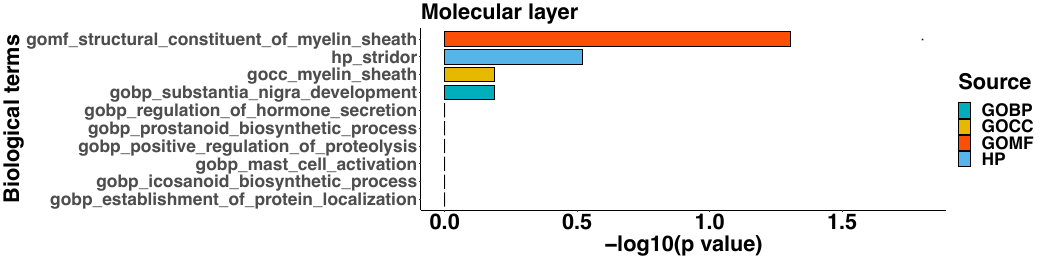 | 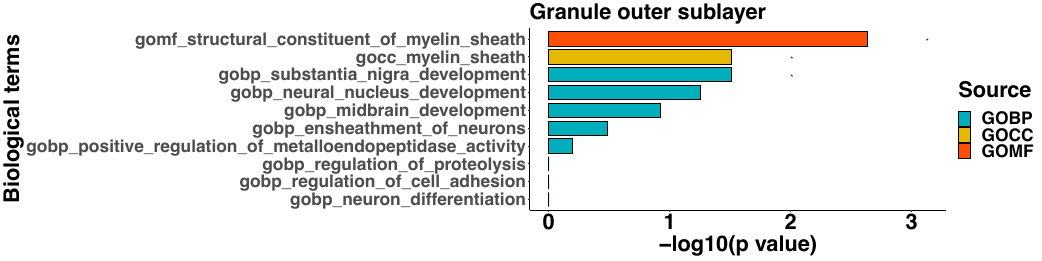 |

**Figure S27.** **Gene set enrichment analysis on the region-specific genes in the Slide-seq data**. (**A**) The top 10 enriched gene sets are shown for each of the eight detected tissue regions. Color represents different data sources for annotating the gene sets.

Figure S28

| **CA1**  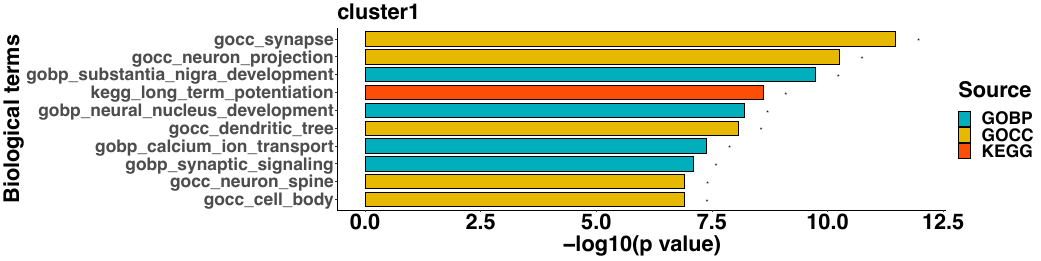 | **Dentate gyrus**  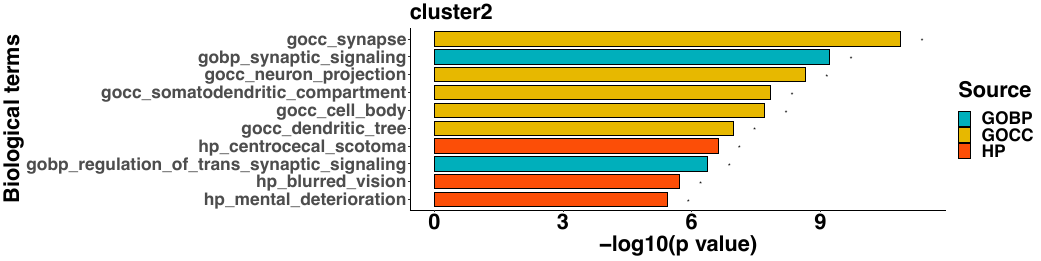 |
| --- | --- |
| **Third ventricle**  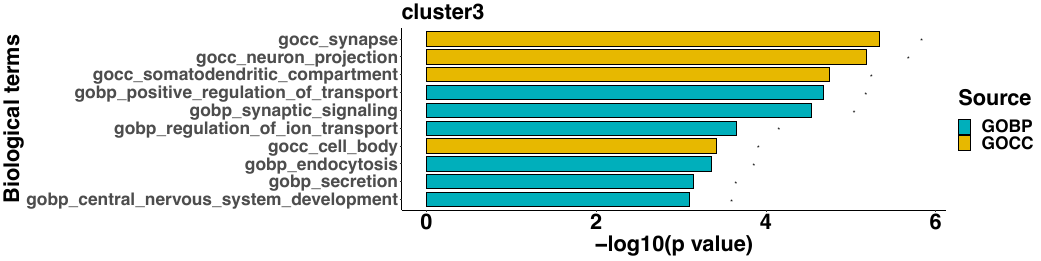 | **CA3**  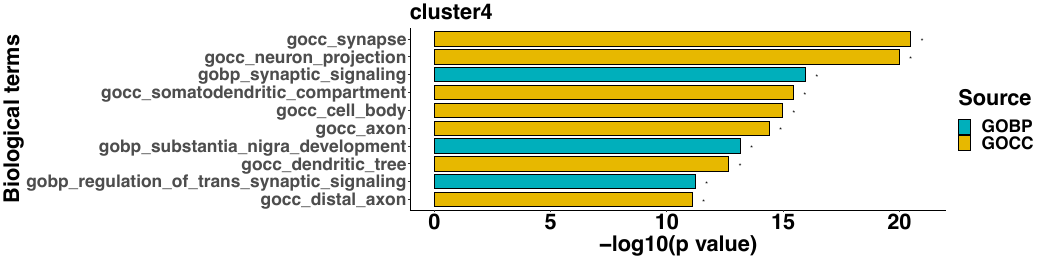 |
| **Layer 6**  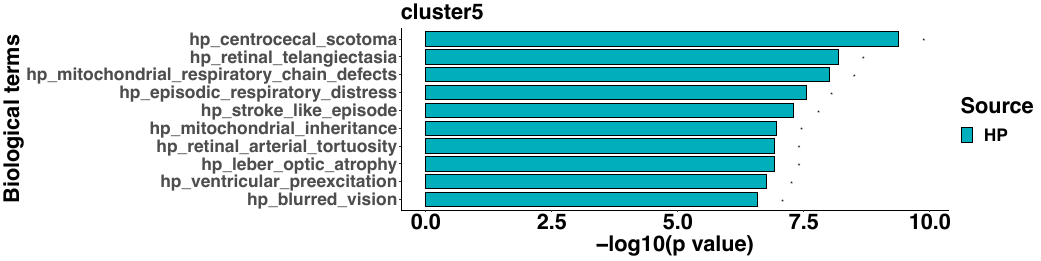 | **Corpus callosum**  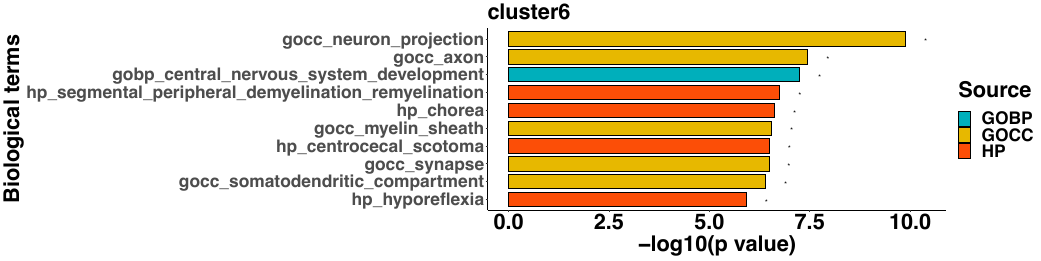 |
| **Hippocampus (slm)**  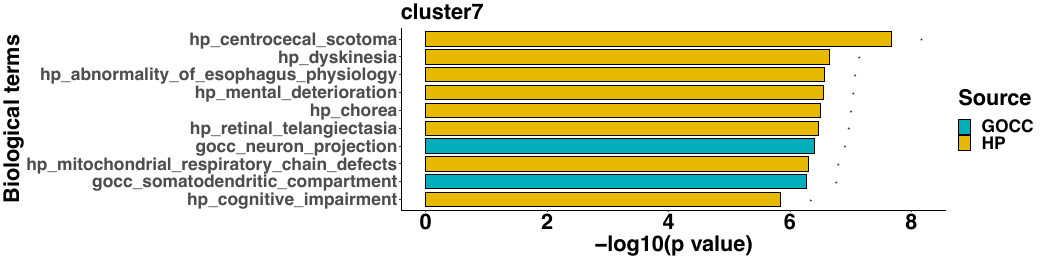 | **Thalamus subregion1**  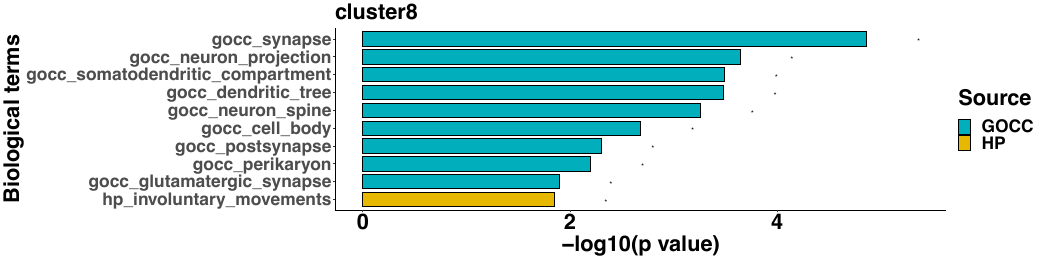 |
| **Layer 4**  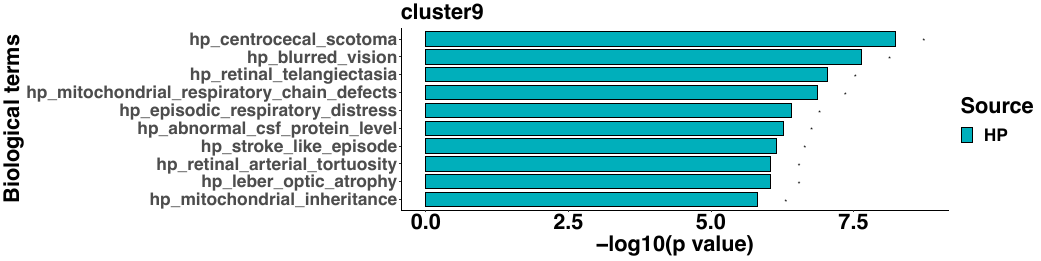 | **Thalamus subregion 3**  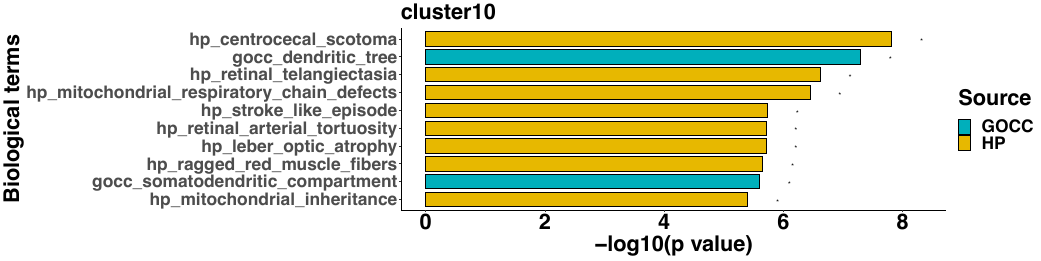 |
| **Hippocampus (so/sr)**  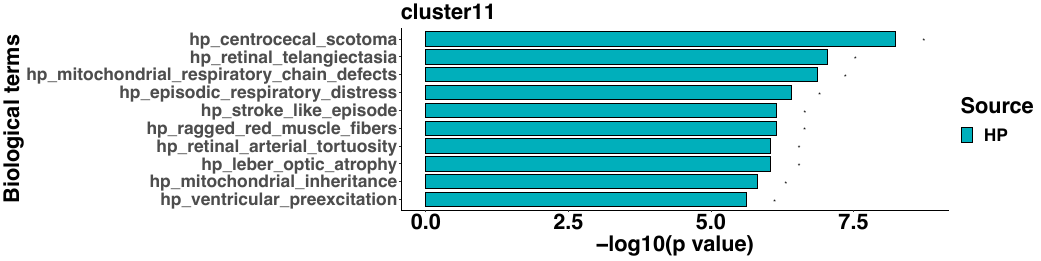 | **Thalamus subregion 2**  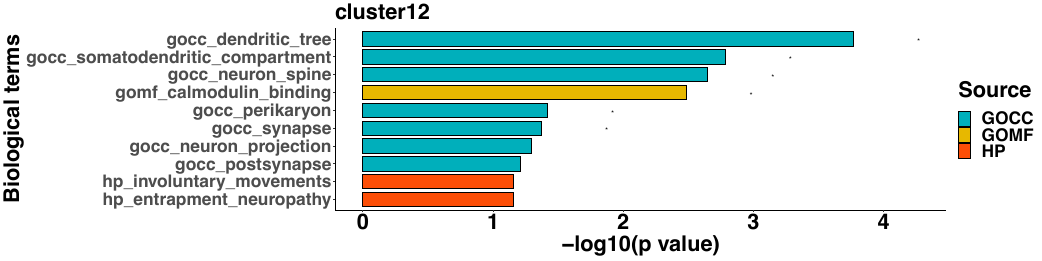 |
| **Layer 5**  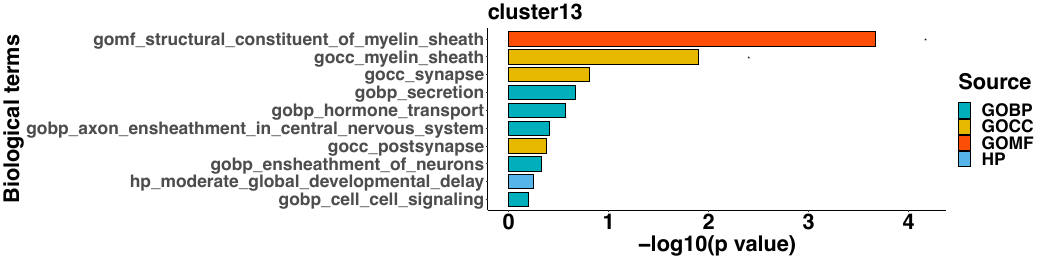 | **Hippocampus (so)**  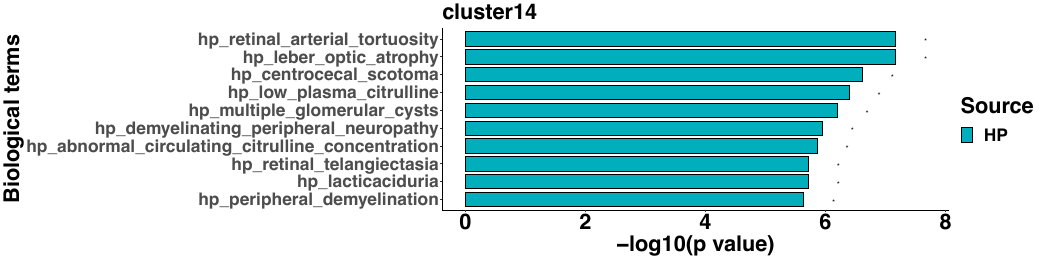 |

**Figure S28.** **Gene set enrichment analysis on the region-specific genes in the Slide-seq V2 data**. (**A**) The top 10 enriched gene sets are shown for each of the eight detected tissue regions. Color represents different data sources for annotating the gene sets.

Figure S29

| 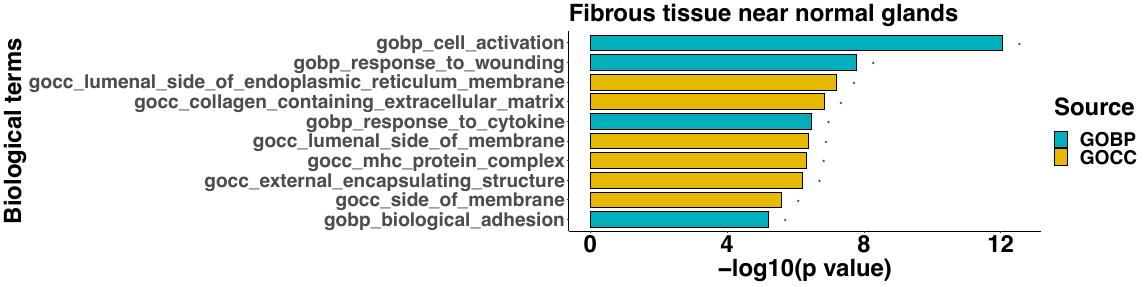 | 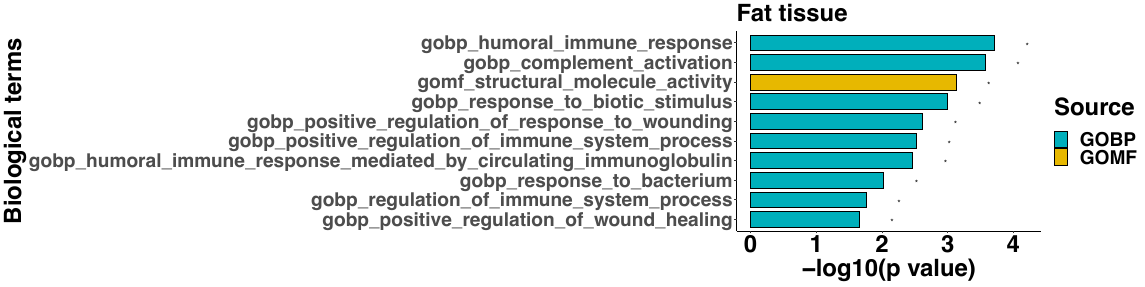 |
| --- | --- |
| 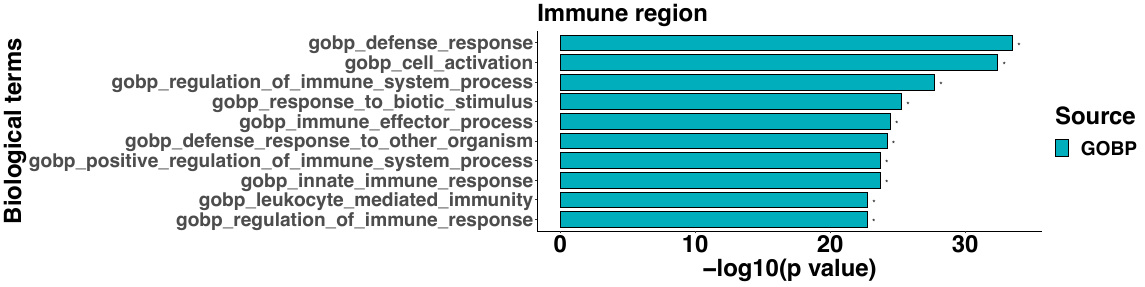 | 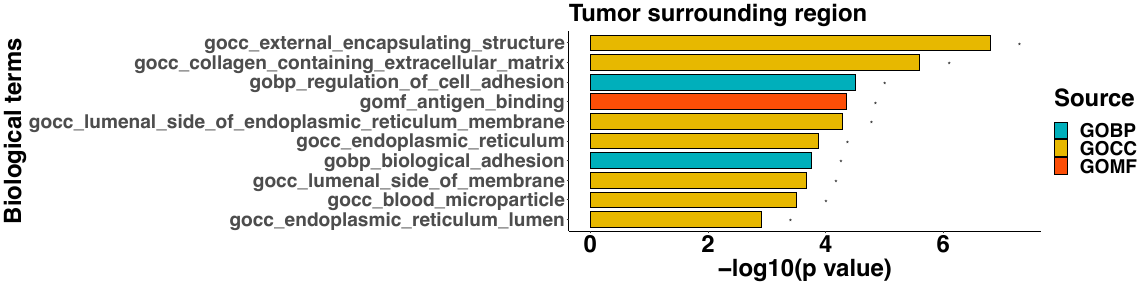 |
| 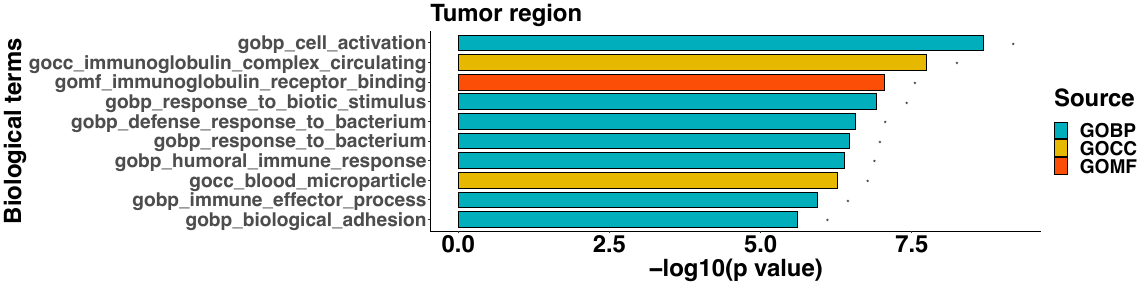 | 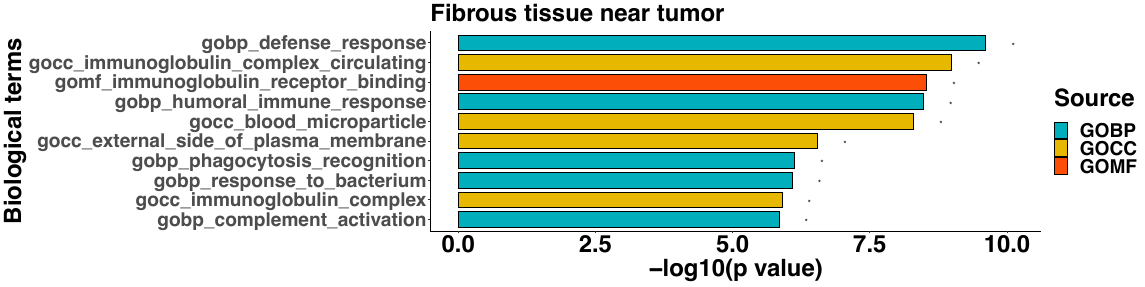 |
| 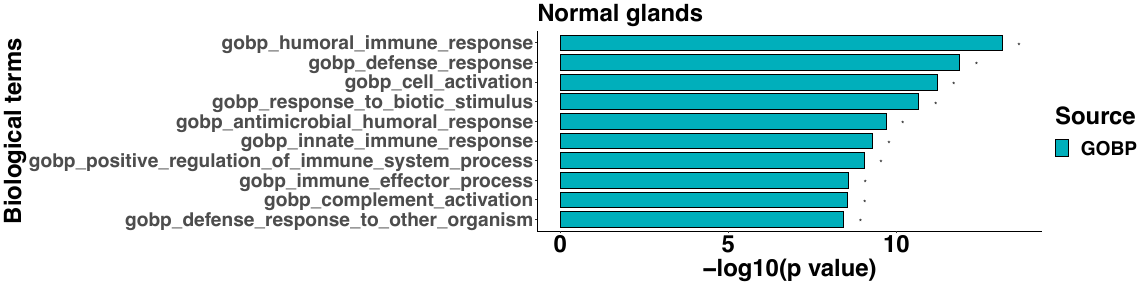 |  |

**Figure S29.** **Gene set enrichment analysis on the region-specific genes in the ST tumor data**. (**A**) The top 10 enriched gene sets are shown for each of the eight detected tissue regions. Color represents different data sources for annotating the gene sets.

Figure S30

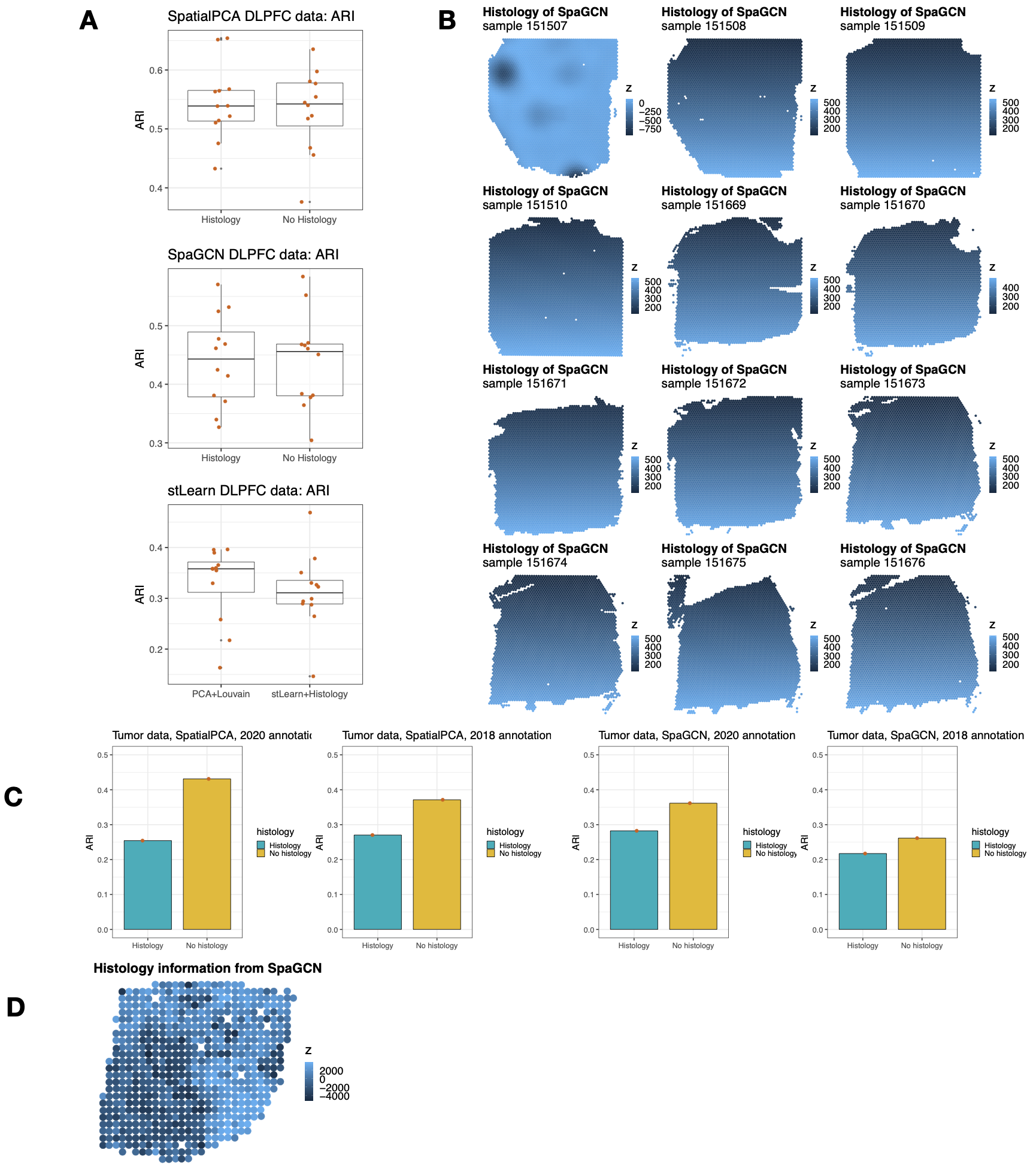

**Figure S30. Clustering results obtained with and without histology information in different methods in DLPFC data and ST tumor data.** (**A**) Clustering results with and without histology information in SpatialPCA, SpaGCN, and stLearn in DLPFC data. The ARI in stLearn was calculated based on SVGs. (**B**) Visualization of the histology information extracted from the RGB values of the H&E image in DLPFC data through SpaGCN. (**C**) Clustering results with and without histology information in SpatialPCA, SpaGCN, and stLearn in the ST tumor data. The ground truth annotations are from the ST tumor data original paper (Andersson et al. 2020) and a earlier version of the paper (Salmén et al. 2018). (**D**) Visualization of the histology information extracted from the RGB values of the H&E image in ST tumor data through SpaGCN.

Figure S31

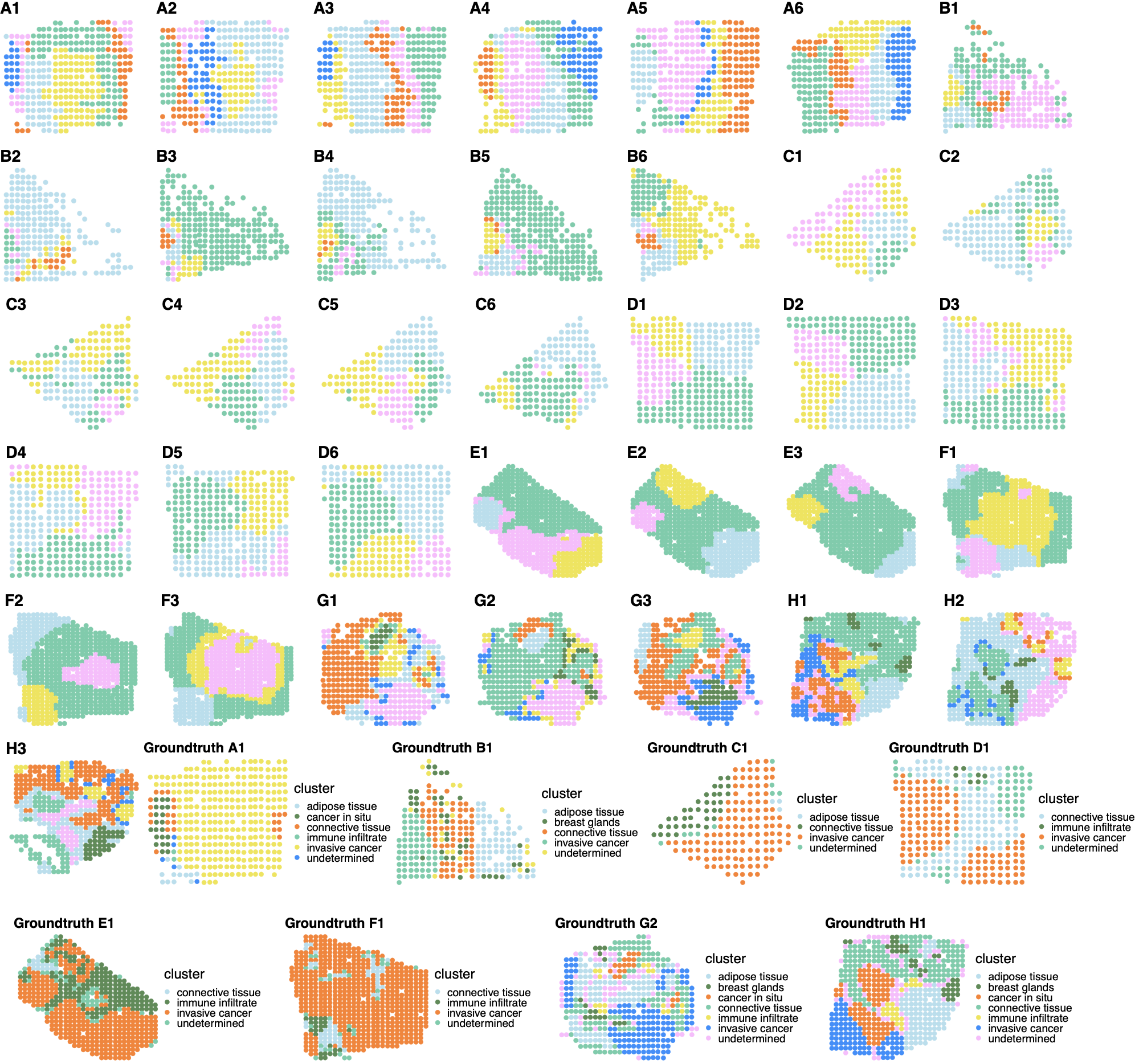

**Figure S31. Clustering results of different spatial domain detection methods in ST tumor across all 36 samples.** **First five rows**: SpatialPCA clustering results of the 36 samples in ST tumor data. **Sixth to seventh rows**: Ground truth annotation of the available A1, B1, C1, D1, E1, F1, G2, and H1 samples**.**

Figure S32

A. B.

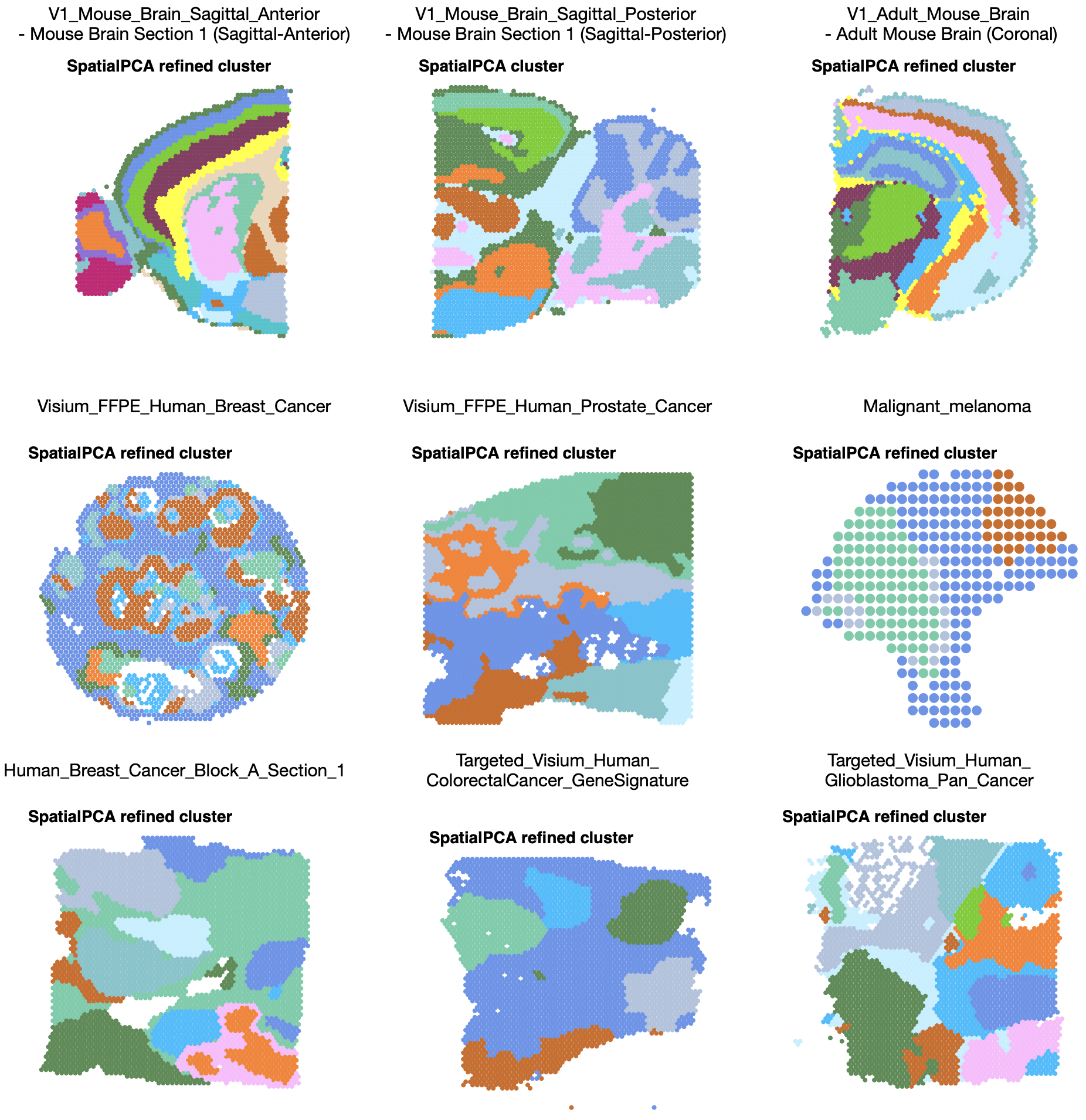

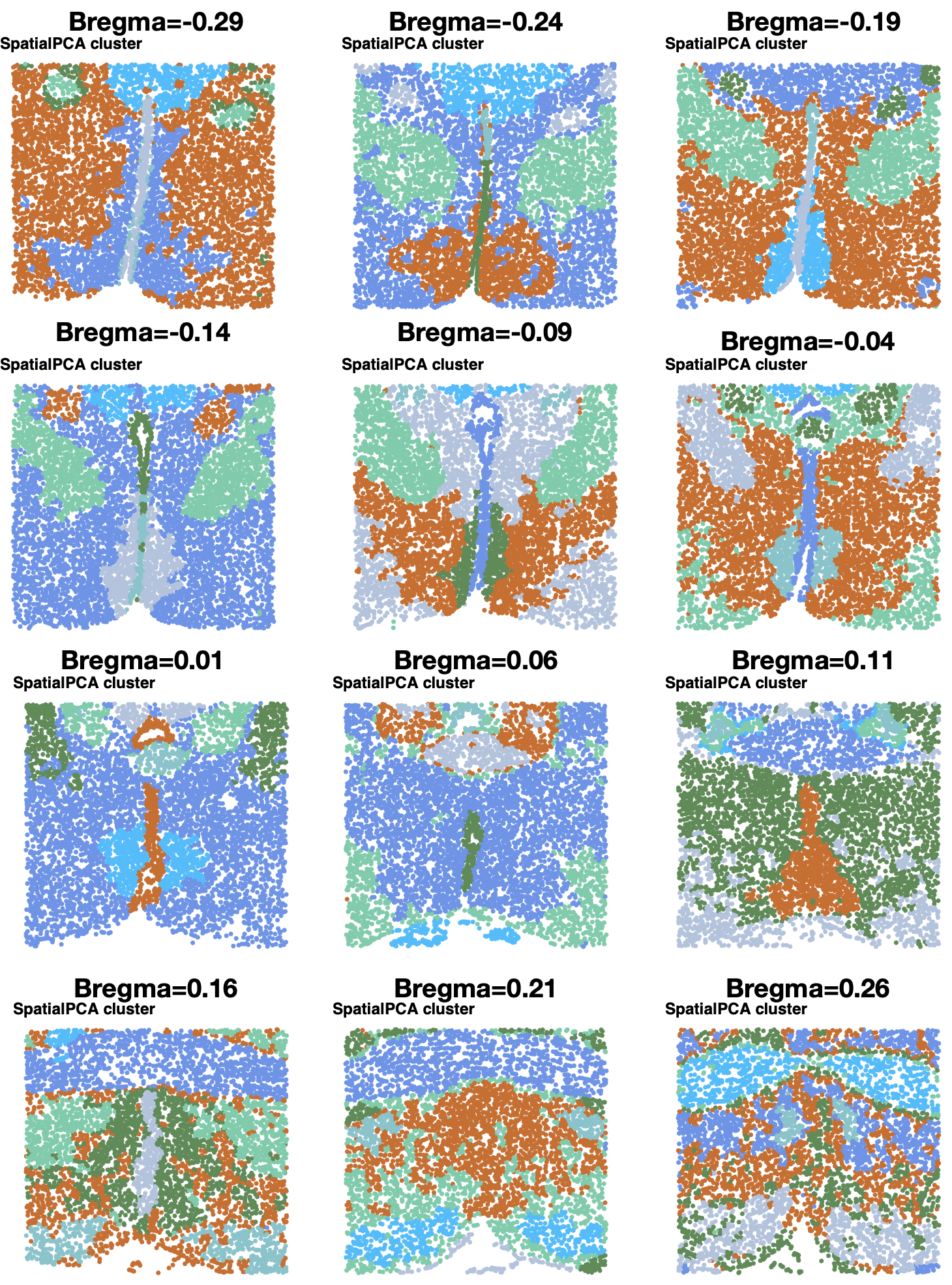

**Figure S32. Clustering results of different spatial domain detection methods in more dataset examples.** (**A**). SpatialPCA clustering results in 10X Visium datasets downloaded from 10X genomics. (**B**) SpatialPCA clustering results in 12 MERFISH samples at bregma values ranging from -0.29 to 0.26.

**Table S1. Median ARI values of simulations at single cell level.** (Default settings in each method are shaded in green color.)

|  | **scenario1** | **scenario2** | **scenario3** | **scenario4** |
| --- | --- | --- | --- | --- |
| **SpatialPCA SVGs** | 0.975 | 0.877 | 0.931 | 0.439 |
| SpatialPCA HVGs | 0.953 | 0.864 | 0.882 | 0.504 |
| SpatialPCA all genes | 0.948 | 0.851 | 0.877 | 0.518 |
| BayesSpace SVGs | 0.56 | 0.08 | 0.205 | 0.005 |
| **BayesSpace HVGs** | 0.646 | 0.225 | 0.286 | 0.075 |
| BayesSpace all genes | 0.641 | 0.167 | 0.215 | 0.064 |
| SpaGCN SVGs | 0.885 | 0.264 | 0.411 | NA |
| SpaGCN HVGs | 0.869 | 0.278 | 0.408 | 0.091 |
| **SpaGCN all genes** | 0.883 | 0.277 | 0.412 | 0.091 |
| PCA SVGs | 0.644 | 0.109 | 0.264 | 0.0002 |
| PCA HVGs | 0.641 | 0.221 | 0.283 | 0.075 |
| PCA all genes | 0.646 | 0.225 | 0.286 | 0.075 |
| NMF SVGs | 0.634 | 0.195 | 0.285 | 0.006 |
| NMF HVGs | 0.593 | 0.205 | 0.266 | 0.069 |
| NMF all genes | 0.572 | 0.201 | 0.254 | 0.068 |

**Table S2. Median ARI values of simulations at spot level (spot diameter is 90um).**

|  | **scenario1** | **scenario2** | **scenario3** | **scenario4** |
| --- | --- | --- | --- | --- |
| SpatialPCA | 0.956 | 0.890 | 0.919 | 0.804 |
| BayesSpace | 0.807 | 0.432 | 0.615 | 0.702 |
| SpaGCN | 0.915 | 0.214 | 0.292 | 0.005 |
| PCA | 0.671 | 0.229 | 0.298 | 0.078 |
| NMF | 0.427 | 0.115 | 0.151 | 0.015 |

**Table S3. Median LISI values of simulations at spot level (spot diameter is 90um).**

|  | **scenario1** | **scenario2** | **scenario3** | **scenario4** |
| --- | --- | --- | --- | --- |
| SpatialPCA | 1.002 | 1.012 | 1.009 | 1.036 |
| BayesSpace | 1.171 | 1.899 | 1.442 | 1.012 |
| SpaGCN | 1.041 | 2.024 | 1.937 | 1.894 |
| PCA | 1.365 | 2.345 | 2.152 | 2.995 |
| NMF | 1.821 | 2.971 | 2.406 | 3.312 |

**Table S4. Median CHAOS values of simulations at spot level (spot diameter is 90um).**

|  | **scenario1** | **scenario2** | **scenario3** | **scenario4** |
| --- | --- | --- | --- | --- |
| SpatialPCA | 0.042 | 0.043 | 0.042 | 0.042 |
| BayesSpace | 0.050 | 0.047 | 0.048 | 0.044 |
| SpaGCN | 0.043 | 0.048 | 0.047 | 0.048 |
| PCA | 0.052 | 0.052 | 0.052 | 0.054 |
| NMF | 0.052 | 0.054 | 0.054 | 0.055 |

**Table S5. Median PAS values of simulations at spot level (spot diameter is 90um).**

|  | **scenario1** | **scenario2** | **scenario3** | **scenario4** |
| --- | --- | --- | --- | --- |
| SpatialPCA | 0.006 | 0.034 | 0.023 | 0.038 |
| BayesSpace | 0.087 | 0.246 | 0.180 | 0.038 |
| SpaGCN | 0.034 | 0.386 | 0.374 | 0.335 |
| PCA | 0.151 | 0.555 | 0.458 | 0.795 |
| NMF | 0.311 | 0.689 | 0.609 | 0.814 |

**Table S6. Median ARI values of 12 samples in DLPFC data.** (Default settings in each method are shaded in green color.)

| **Methods/Gene type** | **SVGs** | **HVGs** | **All genes** |
| --- | --- | --- | --- |
| **SpatialPCA** | **0.542** | 0.403 | 0.303 |
| **BayesSpace** | 0.479 | **0.438** | 0.449 |
| **SpaGCN** | 0.190 | 0.208 | **0.443** |
| **stLearn** (default with marker genes, median ARI=0.470) | 0.311 | 0.342 | 0.345 |
| **PCA** | 0.358 | 0.305 | 0.345 |
| **NMF** | 0.366 | 0.258 | 0.318 |

**Table S7. Number of pseudo-time associated genes detected for each of the trajectory in the DLPFC data.**

| **Dataset** | **Trajectory** **number** | **Number of pseudo-time associated genes** |
| --- | --- | --- |
| DLPFC | 1 | 2763 |
| Slide-seq V2 cortical layer | 1 | 883 |
| ST tumor | 1 | 1713 |

**Table S8. Number of region-specific** **genes in each of the seven spatial domains in the DLPFC data.** The clusters correspond to the region labeling in Figure S26B.

| **Spatial domain name** | **Number of region-specific genes** |
| --- | --- |
| Cluster 1 | 252 |
| Cluster 2 | 265 |
| Cluster 3 | 1327 |
| Cluster 4 | 172 |
| Cluster 5 | 395 |
| Cluster 6 | 68 |
| Cluster 7 | 196 |

**Table S9. Number of region-specific** **genes in each of the eight spatial domains in the Slide-seq data.**

| **Spatial domain name** | **Number of region-specific genes** |
| --- | --- |
| Choroid plexus | 3 |
| White matter | 9 |
| GCL middle layer | 7 |
| GCL inner sublayer | 2 |
| Cerebellum nuclei | 4 |
| Molecular layer | 9 |
| GCL outer layer | 3 |
| Purkinje layer | 14 |

**Table S10. Number of region-specific** **genes in each of the 14 spatial domains in the Slide-seq V2 data.**

| **Spatial domain name** | **Number of region-specific genes** | **Spatial domain name** | **Number of region-specific genes** |
| --- | --- | --- | --- |
| CA1 | 72 | Dentate gyrus | 61 |
| Third ventricle | 34 | CA3 | 114 |
| Layer 6 | 20 | Corpus callosum | 66 |
| Hippocampus (slm) | 39 | Thalamus subregion1 | 24 |
| Layer 4 | 32 | Thalamus subregion 3 | 38 |
| Hippocampus (so/sr) | 32 | Thalamus subregion 2 | 16 |
| Layer 5 | 14 | Hippocampus (so) | 17 |

**Table S11. Number of region-specific** **genes detected in each of the seven spatial domains in the ST tumor data.**

| **Spatial domain name** | **Number of region-specific genes** |
| --- | --- |
| Fibrous tissue near normal glands | 143 |
| Fat tissue | 104 |
| Immune region | 321 |
| Tumor surrounding region | 131 |
| Tumor region | 389 |
| Fibrous tissue near tumor | 81 |
| Normal glands | 369 |

**Table S12. Computation time and peak memory usage for SpatialPCA, BayesSpace, SpaGCN, PCA, and NMF in the three real data applications.** Computing time is recorded in dataset using a single thread on an Intel(R) Xeon(R) Gold 6138 CPU @ 2.00GHz processor.

|  | **DLPFC data (n=3639)** | **Slide-seq data**  **(n=25,551)** | **Slide-seq V2 data (n=53,208)** | **ST data**  **(n=613)** |
| --- | --- | --- | --- | --- |
| **SpatialPCA**  Time / Memory | 12min / 2748Mb | 23min / 11Gb | 6.1 hours / 70Gb | 11s / 157 Mb,  high resolution: 1s (322Mb) |
| **BayesSpace**  Time / Memory | 69min / 8717Mb | 4.1 hours / 40Gb | >100Gb, did not run | 85s / 713Mb,  high resolution: 22min / 1026Mb |
| **SpaGCN**  Time / Memory | 1min / 1092Mb | 6min / 7Gb | 1.2 hours / 45Gb | 11s / 89Mb |
| **PCA**  Time / Memory | 0.5min / 609Mb | 10s / 401Mb | 4min / 3.8Gb | 1s / 21Mb |
| **NMF**  Time / Memory | 0.5min / 870Mb | 37s / 1.2Gb | 5.3min / 9.9Gb | 3s / 51Mb |
| **stLearn**  Time / Memory | 125min / 202Mb | - | - | - |
