## Supplementary Text for "Spatially Aware Dimension Reduction for Spatial Transcriptomics"

**Algorithm for SpatialPCA**

We describe the detailed algorithm for SpatialPCA. Specifically, we first integrate out both $\boldsymbol{B}$ and $\boldsymbol{Z}$ to obtain a marginal likelihood, based on which we infer $\tau, \sigma_{0}^{2}$ and $\boldsymbol{W}$. We then estimate $\boldsymbol{Z}$ by computing their posterior mean conditional on the estimated $\tau, \sigma_{0}^{2}$ and $\boldsymbol{W}$.

We first integrate out $\boldsymbol{B}$. To simplify notation, we denote $\boldsymbol{M=}\boldsymbol{I}_{n}\boldsymbol{-X}\left( \boldsymbol{X}^{T}\boldsymbol{X} \right)^{\boldsymbol{-1}}\boldsymbol{X}^{T}$ and $\boldsymbol{Y}^{*}=\left( \boldsymbol{Y}-\boldsymbol{WZ} \right)^{T}$, where we have $Y_{i}^{*}\sim MVN(X_{i}\boldsymbol{B}, \sigma_{0}^{2}\boldsymbol{I}_{n})$. The marginal distribution for $\boldsymbol{Y}^{*}$ after integrating out ***B*** is:

$$p\left( \boldsymbol{Y}^{\boldsymbol{*}}\mid\sigma_{0}^{2} \right)=\int_{\boldsymbol{B}} L\left( \boldsymbol{B},\sigma_{0}^{2} \right)d\boldsymbol{B}=\int_{\boldsymbol{B}} \frac{1}{\left( 2\pi\sigma_{0}^{2} \right)^{\frac{n}{2}}}e^{-\frac{1}{2\sigma_{0}^{2}}\left( \boldsymbol{Y}^{*}-\boldsymbol{XB} \right)^{T}\left( \boldsymbol{Y}^{*}-\boldsymbol{XB} \right)}d\boldsymbol{B}=\int_{\boldsymbol{B}} \frac{1}{\left( 2\pi\sigma_{0}^{2} \right)^{\frac{mn}{2}}}e^{-\frac{1}{2\sigma_{0}^{2}}\left( \boldsymbol{B}^{T}\boldsymbol{X}^{T}\boldsymbol{XB}-2\boldsymbol{B}^{T}\boldsymbol{X}^{T}\boldsymbol{Y}^{*}+{\boldsymbol{Y}^{*}}^{T}\boldsymbol{Y}^{*} \right)}d\boldsymbol{B}=\int_{\boldsymbol{B}} \frac{1}{\left( 2\pi\sigma_{0}^{2} \right)^{\frac{mn}{2}}}e^{-\frac{1}{2\sigma_{0}^{2}}\left\{ \left( \boldsymbol{B}-\hat{\boldsymbol{B}} \right)^{T}\boldsymbol{X}^{T}\boldsymbol{X}\left( \boldsymbol{B}-\hat{\boldsymbol{B}} \right)+{\boldsymbol{Y}^{*}}^{T}\boldsymbol{Y}^{*}-{\hat{\boldsymbol{B}}}^{T}\boldsymbol{X}^{T}\boldsymbol{X}\hat{\boldsymbol{B}} \right\}}d\boldsymbol{B}=\frac{1}{\left( 2\pi\sigma_{0}^{2} \right)^{\frac{mn}{2}}}\left( 2\pi\sigma_{0}^{2} \right)^{\frac{mq}{2}}\left| \boldsymbol{X}^{T}\boldsymbol{X} \right|^{-\frac{1}{2}}e^{-\frac{1}{2\sigma_{0}^{2}}\left\{ {\boldsymbol{Y}^{*}}^{T}\boldsymbol{Y}^{*}-{\hat{\boldsymbol{B}}}^{T}\boldsymbol{X}^{T}\boldsymbol{X}\hat{\boldsymbol{B}} \right\}}=\left| \boldsymbol{X}^{T}\boldsymbol{X} \right|^{-\frac{1}{2}}\frac{1}{\left( 2\pi\sigma_{0}^{2} \right)^{\frac{m(n-q)}{2}}}e^{-\frac{1}{2\sigma_{0}^{2}}\left\{ {\boldsymbol{Y}^{*}}^{T}\boldsymbol{Y}^{*}-{\hat{\boldsymbol{B}}}^{T}\boldsymbol{X}^{T}\boldsymbol{X}\hat{\boldsymbol{B}} \right\}} (Here \hat{\boldsymbol{B}}=\left( \boldsymbol{X}^{T}\boldsymbol{X} \right)^{-1}\boldsymbol{X}^{T}\boldsymbol{Y}^{*})= \left| \boldsymbol{X}^{T}\boldsymbol{X} \right|^{-\frac{1}{2}}\frac{1}{\left( 2\pi\sigma_{0}^{2} \right)^{\frac{n-q}{2}}}e^{-\frac{1}{2\sigma_{0}^{2}}\left\{ {\boldsymbol{Y}^{*}}^{T}\left( \boldsymbol{I}-\boldsymbol{X}\left( \boldsymbol{X}^{T}\boldsymbol{X} \right)^{-1}\boldsymbol{X}^{T} \right)\boldsymbol{Y}^{*} \right\}}$$

(1)

The marginal distribution for $\boldsymbol{Y}^{*}$ can be simplified as:

$$p\left( \boldsymbol{Y}^{\boldsymbol{*}}\mid\sigma_{0}^{2} \right)\propto\left( \sigma_{0}^{2} \right)^{-\frac{m\left( n-q \right)}{2}}\exp(tr \left( -\frac{{\boldsymbol{Y}^{*}}^{T}\boldsymbol{M}\boldsymbol{Y}^{\boldsymbol{*}}}{2\sigma_{0}^{2}} \right)).$$

(2)

The distribution for $\boldsymbol{Y}$, conditional on $\boldsymbol{Z}$, thus becomes:

$$p\left( \boldsymbol{Y}\mid\boldsymbol{Z},\boldsymbol{W},\sigma_{0}^{2},\tau\right)\propto\left( \sigma_{0}^{2} \right)^{-\frac{m\left( n-q \right)}{2}}\exp(tr \left( -\frac{(\boldsymbol{Y}-\boldsymbol{WZ})\boldsymbol{M}(\boldsymbol{Y}-\boldsymbol{WZ})^{T}}{2\sigma_{0}^{2}} \right)).$$

(3)

The joint likelihood of $\boldsymbol{Y},\boldsymbol{Z}$ is

$$\begin{matrix} p\left( \boldsymbol{Y},\boldsymbol{Z}\mid\boldsymbol{W},\sigma_{0}^{2},\tau\right)\propto p\left( \boldsymbol{Y}\mid\boldsymbol{Z},\boldsymbol{W},\sigma_{0}^{2},\tau\right)p\left( \boldsymbol{Z} | \boldsymbol{W},\sigma_{0}^{2},\tau\right) \\ \propto\left( \sigma_{0}^{2} \right)^{-\frac{m\left( n-q \right)}{2}}\prod_{l=1}^{d} \left| \sigma_{0}^{2}\tau\boldsymbol{K} \right|^{-\frac{1}{2}}\exp(tr \left( -\frac{(\boldsymbol{Y}-\boldsymbol{WZ})\boldsymbol{M}(\boldsymbol{Y}-\boldsymbol{WZ})^{T}+ \sum_{l=1}^{d} {\boldsymbol{Z}_{l}\left( \tau\boldsymbol{K} \right)}^{-1}\boldsymbol{Z}_{l}^{T}}{2\sigma_{0}^{2}} \right)) \\ \propto\left( \sigma_{0}^{2} \right)^{-\frac{m\left( n-q \right)}{2}}\prod_{l=1}^{d} \left| \sigma_{0}^{2}\tau\boldsymbol{K} \right|^{-\frac{1}{2}}\exp(tr \left( -\frac{\boldsymbol{YM}\boldsymbol{Y}^{T}}{2\sigma_{0}^{2}} \right)) \\ \times\exp\left\{ -\frac{\boldsymbol{ZM}\boldsymbol{Z}^{T}-2\boldsymbol{Z}^{T}\left( \boldsymbol{W}^{T}\boldsymbol{M} \right)\boldsymbol{Y}+\sum_{l=1}^{d} {\boldsymbol{Z}_{l}\left( \tau\boldsymbol{K} \right)}^{-1}\boldsymbol{Z}_{l}^{T}}{2\sigma_{0}^{2}} \right\}. \end{matrix}$$

(4)

Next, we integrate out $\boldsymbol{Z}$ to obtain the marginal distribution for $\boldsymbol{Y}$:

$$\begin{matrix} & p(\boldsymbol{Y}|\left. \boldsymbol{W},\sigma_{0}^{2},\tau\right) \\ & \propto\left( \sigma_{0}^{2} \right)^{-\frac{m(n-q)}{2}}\prod_{l=1}^{d} \left| \tau\boldsymbol{MK}+\boldsymbol{I}_{n} \right|^{-\frac{1}{2}}\exp(tr\left( -\frac{\boldsymbol{YM}\boldsymbol{Y}^{T}}{2\sigma_{0}^{2}} \right)) \\ & \times\exp\left\{ -\frac{\sum_{l=1}^{d} \boldsymbol{Y}^{T}\left( \boldsymbol{W}^{T}\boldsymbol{M} \right)^{T}\left( \boldsymbol{M}+{\tau^{-1}\boldsymbol{K}}^{-1} \right)^{-1}\left( \boldsymbol{W}^{T}\boldsymbol{M} \right)\boldsymbol{Y}}{2\sigma_{0}^{2}} \right\} \\ & \propto\left( \sigma_{0}^{2} \right)^{-\frac{m(n-q)}{2}}\prod_{l=1}^{d} \left| \tau\boldsymbol{MK}+\boldsymbol{I}_{n} \right|^{-\frac{1}{2}} \\ & \times\exp\left\{ -\frac{\mathrm{tr} \left( \boldsymbol{YMY}^{T} \right)- \sum_{l=1}^{d} \boldsymbol{w}_{l}^{T}\boldsymbol{YM}\left( \boldsymbol{M}+{\tau^{-1}\boldsymbol{K}}^{-1} \right)^{-1}\boldsymbol{M}\boldsymbol{Y}^{T}\boldsymbol{w}_{l}}{2\sigma_{0}^{2}} \right\}. \end{matrix}$$

(5)

Based on the marginal distribution of $\boldsymbol{Y}$, we can obtain the maximum likelihood estimator of $\sigma_{0}^{2}$ as

$$\hat{\sigma}_{0}^{2}=\frac{tr\left( \boldsymbol{YMY}^{T} \right)-\sum_{l=1}^{d} \boldsymbol{w}_{l}^{T}\boldsymbol{YM}\left( \boldsymbol{M}+{\tau^{-1}\boldsymbol{K}}^{-1} \right)^{-1}\boldsymbol{M}\boldsymbol{Y}^{T}\boldsymbol{w}_{l}}{m(n-q)}.$$

(6)

Above, $\boldsymbol{w}_{l}$ is the *l-*th column of the loading matrix $\boldsymbol{W}$.

Denote$S=tr\left( \boldsymbol{YMY}^{T} \right)-\sum_{l=1}^{d} \boldsymbol{w}_{l}^{T}\boldsymbol{YM}\left( \boldsymbol{M}+{\tau^{-1}\boldsymbol{K}}^{-1} \right)^{-1}\boldsymbol{M}\boldsymbol{Y}^{T}\boldsymbol{w}_{l}$, plugging the maximum likelihood estimator of $\sigma_{0}^{2}$ back to the marginal distribution of $\boldsymbol{Y}$ in equation (5) gives

$$\begin{matrix} & p(\boldsymbol{Y}|\left. \boldsymbol{W},\hat{\sigma}_{0}^{2},\tau\right) \\ & \propto{|S|}^{-\frac{m(n-q)}{2}}\prod_{l=1}^{d} \{{\left| \tau\boldsymbol{MK}+\boldsymbol{I}_{n} \right|\}}^{-\frac{1}{2}} \\ & \propto{|S|}^{-\frac{m(n-q)}{2}}\prod_{l=1}^{d}\left\{ \left| \tau\boldsymbol{K}+\boldsymbol{I}_{n}||\boldsymbol{I}_{n}-\left( \tau\boldsymbol{K}+\boldsymbol{I}_{n} \right)^{-1}\tau\boldsymbol{KX}\left( \boldsymbol{X}^{T}\boldsymbol{X} \right)^{-1}\boldsymbol{X}^{T} \right| \right\}^{-\frac{1}{2}} \\ & \propto{|S|}^{-\frac{m(n-q)}{2}}\prod_{l=1}^{d} \{{\left| \tau\boldsymbol{K}+\boldsymbol{I}_{n}||\boldsymbol{X}^{T}\boldsymbol{X} \right|^{-1}\left| \boldsymbol{X}^{T}\left( \tau\boldsymbol{K}+\boldsymbol{I}_{n} \right)^{-1}\boldsymbol{X} \right|\}}^{-\frac{1}{2}} \\ & \propto{|S|}^{-\frac{m(n-q)}{2}}\prod_{l=1}^{d}\{ \left| \tau\boldsymbol{K}+\boldsymbol{I}_{n} \right|^{-\frac{1}{2}}\left| \boldsymbol{X}^{T}\left( \tau\boldsymbol{K}+\boldsymbol{I}_{n} \right)^{-1}\boldsymbol{X} \right|^{-\frac{1}{2}}\}. \end{matrix}$$

(7)

We further maximize the above marginal distribution to obtain the estimators for $\boldsymbol{W}$ and $\tau$:

$$\hat{\boldsymbol{W}}=\underset{\boldsymbol{W}}{argmax}\sum_{l=1}^{d} \boldsymbol{w}_{l}^{T}\boldsymbol{YM}\left( \boldsymbol{M}+{\tau^{-1}\boldsymbol{K}}^{-1} \right)^{-1}\boldsymbol{M}\boldsymbol{Y}^{T}\boldsymbol{w}_{l}, \text{ s.t. } \boldsymbol{W}^{T}\boldsymbol{W}=\boldsymbol{I}_{d},$$

(8)

$$\begin{matrix} \hat{\tau} & =\underset{\tau}{argmax}p(\boldsymbol{Y}\mid\hat{\boldsymbol{W}},\tau). \end{matrix}$$

(9)

We use the Brent’s optimization method implemented in the *optim* function in R to obtain the estimation of $\tau$ in equation (9). We obtain the closed-form expression^1; 2^ of $\hat{\boldsymbol{W}}$ in the form of $\hat{\boldsymbol{W}}=\boldsymbol{LR}\mathbf{,}$where $\boldsymbol{L}$is a *m* by *d* matrix for the first *d* eigenvectors of $\boldsymbol{YM}\left( \boldsymbol{M}+{\tau^{-1}\boldsymbol{K}}^{-1} \right)^{-1}\boldsymbol{M}\boldsymbol{Y}^{T}$and $\boldsymbol{R}$is an arbitrary *d* by *d* orthogonal rotation matrix.

In addition, based on equation (4), we can obtain the maximum likelihood estimate for each $\boldsymbol{Z}_{l}$ as

$$\boldsymbol{Z}_{l}|\boldsymbol{Y},\hat{\boldsymbol{W}},\hat{\sigma}_{0}^{2},\hat{\tau} \sim MVN\left( {\hat{\boldsymbol{Z}}}_{l},{\hat{\boldsymbol{\Sigma}}}_{Z_{l}} \right),$$

(10)

where ${\hat{\boldsymbol{Z}}}_{l}\boldsymbol{=}\left( \boldsymbol{M+}\hat{\tau}^{-1}\boldsymbol{K}^{-1} \right)^{-1}\boldsymbol{M}\boldsymbol{Y}^{T}\hat{\boldsymbol{w}_{l}}$, and ${\hat{\boldsymbol{\Sigma}}}_{Z_{l}}=\hat{\sigma}_{0}^{2}\left( \boldsymbol{M+}\hat{\tau}^{-1}\boldsymbol{K}^{-1} \right)^{-1}$. From equation (1) we can further obtain $\hat{\boldsymbol{B}}=\left( \boldsymbol{X}^{T}\boldsymbol{X} \right)^{-1}\boldsymbol{X}^{T}(\boldsymbol{Y}^{T}-{\hat{\boldsymbol{Z}}}^{T}{\hat{\boldsymbol{W}}}^{T})$.

Carrying out the above inference algorithm requires calculating the following three quantities: $\left( \boldsymbol{M}+\tau^{-1}\boldsymbol{K}^{-1} \right)^{-1}$ in equations (6), (8) and (10), as well as $|\tau\boldsymbol{K}+\boldsymbol{I}_{n}|$ and $\boldsymbol{|X}^{T}\left( \tau\boldsymbol{K}+\boldsymbol{I}_{n} \right)^{-1}\boldsymbol{X|}$ in equation (7). Each quantity in equations (6)-(8) needs to be re-evaluated in each iteration of the inference algorithm as $\tau$ is being updated while the quantity in equation (10) needs to be evaluated once at the last iteration. To improve the computation efficiency of the above inference algorithm, we first perform eigen decomposition on the kernel matrix $\boldsymbol{K=UD}\boldsymbol{U}^{T}$ at the beginning of the optimization, where $\mathbf{D}=diag(\delta_{1}, \ldots, \delta_{n})$ with $\delta_{i}$ being the eigen values, and $\boldsymbol{U}$ is the eigenvector matrix. With the eigen decomposition of $\boldsymbol{K}$, we can simplify the calculation of the three quantities.

Specifically, for $\left( \boldsymbol{M}+\tau^{-1}\boldsymbol{K}^{-1} \right)^{-1}$, it can be calculated using the Woodbury formula as:

$$\begin{matrix} \left( \boldsymbol{M}+\tau^{-1}\boldsymbol{K}^{-1} \right)^{-1} \\ =\left( \boldsymbol{I}-\boldsymbol{X}\left( \boldsymbol{X}^{T}\boldsymbol{X} \right)^{-1}\boldsymbol{X}^{T}+\tau^{-1}\boldsymbol{K}^{-1} \right)^{-1} \\ =\left( \boldsymbol{I}+\tau^{-1}\boldsymbol{K}^{-1}+\boldsymbol{X}\left( -\left( \boldsymbol{X}^{T}\boldsymbol{X} \right)^{-1} \right)\boldsymbol{X}^{T} \right)^{-1} \\ =\left( \boldsymbol{I}+\tau^{-1}\boldsymbol{K}^{-1} \right)^{-1}-\left( \boldsymbol{I}+\tau^{-1}\boldsymbol{K}^{-1} \right)^{-1}\boldsymbol{X}\left[ -\boldsymbol{X}^{T}\boldsymbol{X}+\boldsymbol{X}^{T}\left( \boldsymbol{I}+\tau^{-1}\boldsymbol{K}^{-1} \right)^{-1}\boldsymbol{X} \right]^{-1}\boldsymbol{X}^{T}\left( \boldsymbol{I}+\tau^{-1}\boldsymbol{K}^{-1} \right)^{-1}. \end{matrix}$$

where

$$\begin{matrix} \left( \boldsymbol{I}+\tau^{-1}\boldsymbol{K}^{-1} \right)^{-1} \\ =\left( \boldsymbol{I}+\boldsymbol{U}\left( \tau^{-1}\boldsymbol{D}^{-1} \right)\boldsymbol{U}^{T} \right)^{-1} \\ ={\boldsymbol{U}\left( \boldsymbol{I}+\tau^{-1}\boldsymbol{D}^{-1} \right)}^{-1}\boldsymbol{U}^{T} \end{matrix}$$

(11)

Therefore, we can compute $\boldsymbol{U}^{T}\boldsymbol{M}\boldsymbol{Y}^{T}$at the beginning of the algorithm and then evaluate the quantities in equations (6) and (8) in each iteration of algorithm with a linear complexity with respect to *n*. In addition, we can evaluate $\left( \boldsymbol{M}+\tau^{-1}\boldsymbol{K}^{-1} \right)^{-1}\boldsymbol{M}\boldsymbol{Y}^{T}$in equation (10) in the last iteration with a quadratic complexity with respect to *n*.

For $|\tau\boldsymbol{K}+\boldsymbol{I}_{n}|$, we can express it as $|\tau\boldsymbol{K}+\boldsymbol{I}_{n}|\boldsymbol{=}|\tau\boldsymbol{D+}\boldsymbol{I}_{n}|$, which has a linear complexity with respect to *n*. For $\boldsymbol{|X}^{T}\left( \tau\boldsymbol{K}+\boldsymbol{I}_{n} \right)^{-1}\boldsymbol{X}|$, we first calculate $\left( \tau\boldsymbol{K}+\boldsymbol{I}_{n} \right)^{-1}=\boldsymbol{U}\left( \boldsymbol{I}_{n}+\tau\boldsymbol{D} \right)^{-1}\boldsymbol{U}^{T}$ and then calculate $\left( \tau\boldsymbol{K}+\boldsymbol{I}_{n} \right)^{-1}\boldsymbol{X}$ and $\boldsymbol{X}^{T}\left( \tau\boldsymbol{K}+\boldsymbol{I}_{n} \right)^{-1}\boldsymbol{X}$ before taking its determinant. Because we can compute $\boldsymbol{U}^{T}\boldsymbol{X}$ at the beginning of the algorithm, evaluating $\boldsymbol{|X}^{T}\left( \tau\boldsymbol{K}+\boldsymbol{I}_{n} \right)^{-1}\boldsymbol{X}|$ in each iteration has a linear complexity with respect to *n*.

To further facilitate computation, we applied low-rank approximation in the eigen decomposition step for $\boldsymbol{K}$. In particular, we use the “RSpectra” R package in the eigen decomposition to only obtain the top *r* eigenvectors and *r* eigenvalues. Therefore, both $\boldsymbol{U}$ and $\boldsymbol{D}$ become low-rank matrices with dimension *n* by *r* for $\boldsymbol{U}$ and *r* by *r* for $\boldsymbol{D}$. In this case, we calculate $\left( \boldsymbol{I}+\tau^{-1}\boldsymbol{K}^{-1} \right)^{-1}$ in equation (11) as

$$\left( \boldsymbol{I}+\tau^{-1}\boldsymbol{K}^{-1} \right)^{-1}=\boldsymbol{I}-\boldsymbol{U}\left( \tau\boldsymbol{D}+\boldsymbol{U}^{T}\boldsymbol{U} \right)^{-1}\boldsymbol{U}^{T},$$

and evaluate $\hat{\boldsymbol{Z}}$ in equation (10) as

$$\begin{matrix} \hat{\boldsymbol{Z}}\boldsymbol{=}{\hat{\boldsymbol{W}}}^{\boldsymbol{T}}\boldsymbol{YM}\left( \boldsymbol{M}+\tau^{-1}\boldsymbol{K}^{-1} \right)^{-1}=\hat{\tau}{\hat{\boldsymbol{W}}}^{\boldsymbol{T}}\boldsymbol{YMK-}\hat{\tau}{\hat{\boldsymbol{W}}}^{\boldsymbol{T}}\boldsymbol{YM}\boldsymbol{U}\left( \tau^{-1}\boldsymbol{D}^{-1}+\boldsymbol{U}^{T}\boldsymbol{MU} \right)^{-1}\boldsymbol{U}^{T}\boldsymbol{MK.} \end{matrix}$$

In the above low-rank approximation, we choose the rank *r* as a function of sample size. Specifically, for data with a large sample size (n>5,000), we evaluated quantities in equations (6) and (8) in each iteration using a small *r*=20, since these quantities are insensitive to the choice of rank *r*. We obtained the estimates ${\hat{\boldsymbol{Z}}}_{l}$ in equation (10) using a relatively large *r*, with *r* set to be 10% of the sample size *n*, which ensures the top *r* eigen values to explain at least 90% of the variance in the present study. For data with a small sample size (n$\leq$5,000), we use the same *r* to evaluate quantities in equations (6), (8) and (10), with *r* chosen to ensure that the top *r* eigen values explain at least 90% of the variance. Our software implementation also allows users to specify their own choice of *r*.

Overall, the computational time complexity of our algorithm is *O*(*tdm*^2^+*rn*^2^), where *t* represents the number of iterations in the optimization algorithm, with memory requirement being *O*(*mn*+*n*^2^).

**References.**

1. Kokiopoulou, E., Chen, J., and Saad, Y. (2011). Trace optimization and eigenproblems in dimension reduction methods. Numer Linear Algebr 18, 565-602.

2. Saad, Y. (2011). Numerical Methods for Large Eigenvalue Problems Preface to the Classics Edition. Class Appl Math 66, Xiii-+.
